# The Ligand Preference of LRP1 is Regulated by O-glycans

**DOI:** 10.1101/2025.10.23.684103

**Authors:** John Hintze, Asli Bahar Topaktas, Thomas M. Daugbjerg, Shifa Jebari, Silvia D’Andrea, Noortje de Haan, Lasse H. Hansen, Rob Baars, Noé Quittot, Cesar Martin, Zhang Yang, Sergey Y. Vakhrushev, Rebecca L. Miller, Dudley K. Strickland, Bradley Hyman, Elisa Fadda, Katrine T. Schjoldager

## Abstract

Low-density lipoprotein receptor (LDLR) and LDLR-related proteins (LRPs) are endocytic receptors serving as essential physiological regulators of multiple processes including cholesterol clearance, protein reabsorption and neuronal protein trafficking. We originally discovered O-glycans in linkers of the ligand-binding domains of LDLR/LRP receptors and showed that these play critical roles for uptake of LDL by LDLR and albumin by LRP2. Remarkably, these linker O-glycans are introduced exclusively by GALNT11, one out of 20 polypeptide GalNAc-transferase isoenzymes. Here, we investigate the role of linker O-glycans on the large (∼600 kDa) and widely expressed multiligand LRP1 receptor implicated in diseases including neuropathies. In genetically engineered cell models we activated endogenous full-coding LRP1 with and without O-glycans and demonstrate that while the uptake of certain ligands, such as RAP and ApoE, was unaffected by O-glycans, the uptake of the neurotoxic molecules tau and amyloid beta was altered and in opposite directions. This demonstrates that O-glycans in ligand-binding domains can differentially modulate ligand affinity and specificity of LRP1. Our findings highlight an overlooked regulatory mechanism of endocytic receptors and identify the ligand repertoire of LRP1 as being influenced by O-glycans, with potential implications for neurodegenerative disease.

## Introduction

Endocytosis, the process by which cells specifically take up macromolecules from the extracellular environment, is fundamental to all cells. The low-density lipoprotein receptor (LDLR) and LDLR-related proteins (LRPs) comprise a family of structurally related endocytic receptors with essential functions in cholesterol clearance, protein reabsorption from urine and transcytotic trafficking of neuronal and cerebrospinal fluid (CSF) proteins at the blood-CSF and blood-brain-barrier (BBB)^1,2^. Genetic deficiencies in LRP genes are thus associated with a range of diseases including dyslipidemia^3^, Alzheimer’s^4^, migraine^5^ and kidney failure^6^, underscoring their physiological importance.

While LDLR and VLDLR serve a select subset of ligands, mainly LDL and VLDL, respectively, the large (>600 kDa) LRP1 and LRP2/Megalin receptors each serve more than 50 structurally and functionally diverse ligands destined for degradation and/or transcytosis^7–10^. How the broad ligand specificities of LRP1 and LRP2 are accommodated and how different ligands can elicit different cellular responses is unclear, but partly ascribed to the extended and flexible structure of these large receptors with up to four clusters of ligand-binding LDLR class A cysteine-rich complement type repeats (CRs)^11^. Understanding genetic disease susceptibilities and pathologies associated with endocytic receptors^12,13^, requires identification of the ligand repertoire and mechanisms of ligand recognition and selection. Previously we showed that the short linker sequences (**C_6_**X_3-5_**T*C_1_**, *indicates O-glycan at the Thr/Ser residue) in-between CRs often contain an O-glycan that is specifically attached by GALNT11, one of up to 20 distinct polypeptide GalNAc-transferase (GALNT) isoenzymes that initiate protein GalNAc-type O-glycosylation^14–18^. This O-glycan sequence motif is present in CR linkers of LDLR and LRPs and highly conserved in the orthologous genes in vertebrates^15^. Genetic ablation of *GALNT11* resulting in loss of CR O-glycans greatly affected LDLR binding and uptake of LDL^15^ and Lrp2 binding of albumin and kidney function^17^.

LRP1, also known as the Alpha-2-Macroglobulin (A2M) receptor or Apolipoprotein E (ApoE) receptor, is the closest paralog to LRP2 and abundantly expressed in liver, lung and brain and in mice systemic loss of *Lrp1* results in embryonic lethality^19^. LRP1 is synthesized as a ∼600 KDa type 1 transmembrane protein and proteolytically modified by furin in the trans-Golgi network (TGN) to produce an 85 kDa transmembrane subunit (β) and a >500 kDa soluble subunit (α) which remain non-covalently attached as the receptor is presented at the cell surface^9^. The α-subunit contains four ligand-binding CR clusters (I-IV) (**Fig. 1**), harboring in total 31 CRs that serve a diverse group of ligands, including alpha synuclein, amyloid beta (Aβ), and tau. Uptake and transcytosis of ligands by LRP1 drive critical physiological processes related to neurodegenerative disorders^19–22^.

**Fig. 1.**
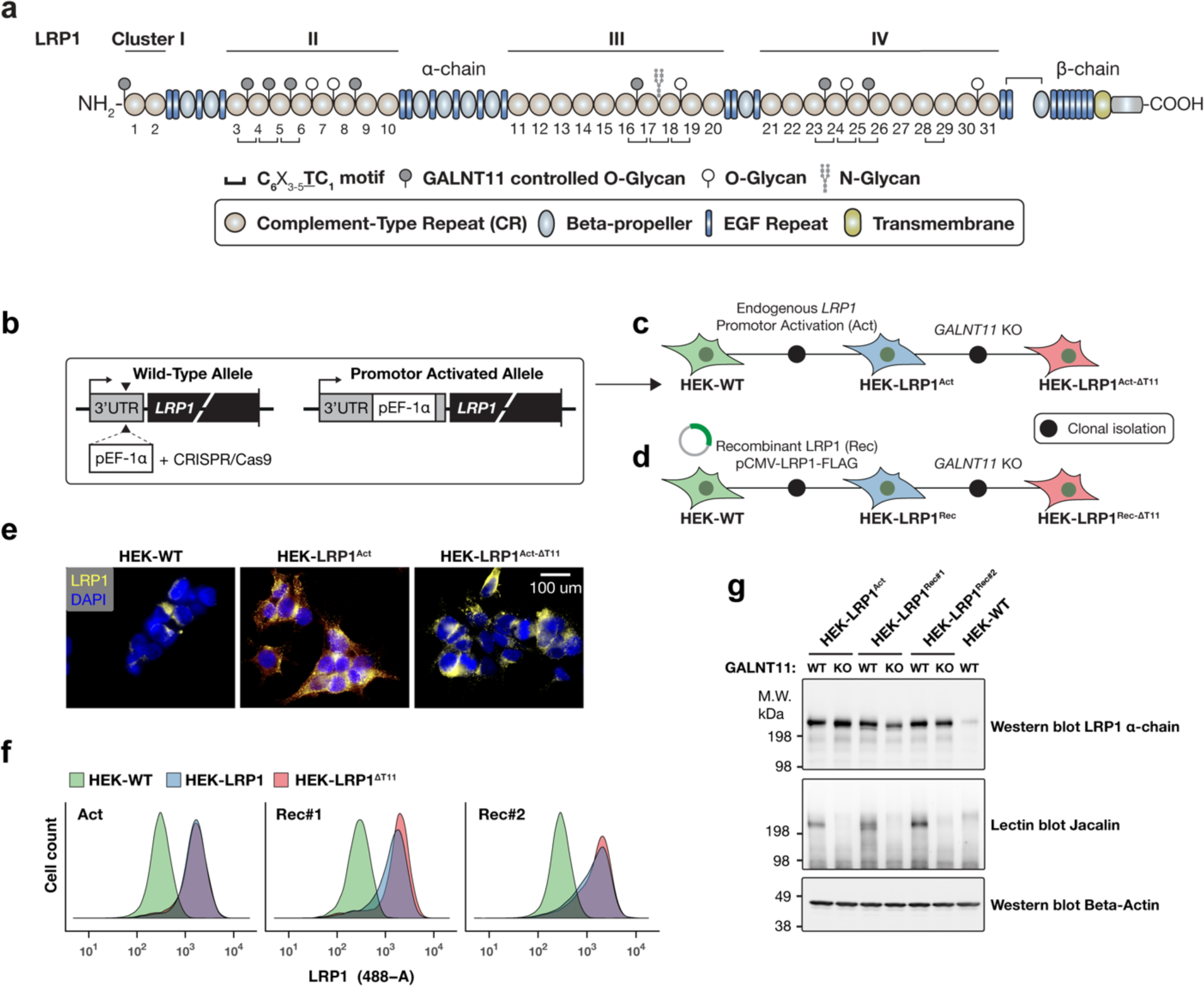
LRP1 CR linkers carries O-glycans. **a** Schematic representation of full-coding LRP1, illustrating the ligand-binding CR domains (numbered 1-31) within clusters I-IV. All adjacent CRs are separated by short linkers and those linker sequences that adhere to the **C**_6_X_3-5_**<u>T</u>C**_1_ motif have been indicated by horizontal brackets. Linker O-glycans identified by O-glycoproteomic analyses of various mammalian tissues and cell lines of wild-type and *GALNT11*^-/-^ genotype are marked by a filled (GALNT11-specific) or open (unkown specificity) circles. **b** Illustration of the promotor activation strategy used to stably induce LRP1 expression by CRISPR/Cas9 mediated knock-in of a dsDNA fragment encoding the EF-1α promoter 140 bp upstream of the endogenous *LRP1* start codon. **c** Lineage ancestry of isogenic HEK cell lines for expression of LRP1 by promotor activation (Act). **d** Lineage ancestry of isogenic HEK cell lines stably transfected with a puromycin selected construct for recombinant expression of full-coding LRP1 with C-terminal 2xFLAG-tag (Rec). Two independent lines (#1 and #2) were selected for *GALNT11* KO by CRISP/Cas9. **e** Immunofluorescence images of fixed HEK-WT, HEK-LRP1^Act^ and HEK-LRP1^Act-ΔT11^ cells stained with anti-LRP1 mAb 8G1. **f** Flow cytometric analysis of cells as indicated stained with anti-LRP1 mAb 8G1. **g** Western and lectin blot analysis of total cell lysates subjected to SDS-PAGE using the anti-LRP1 mAb 8G1 and the Core1/T specific lectin Jacalin. Antibody to β-actin included as loading control.

Here, we aimed to dissect putative functional roles of CR linker O-glycans in LRP1. To this end we established cell models with expression of full-coding LRP1 with and without GALNT11 directed O-glycans, used these to probe uptake of fluorescently labeled ligands and demonstrate that CR linker O-glycans impacted binding and uptake of ligands differently by flow cytometry. While the Receptor-Associated Protein (RAP), A2M and ApoE ligands were impervious to O-glycosylation of LRP1, O-glycosylation significantly enhanced uptake of tau and decreased uptake of Aβ in two cell models, Human Embryonic Kidney 293 (HEK) and neuroblastoma derived SH-SY5Y cells. We further developed a cell-based assay using mass spectrometry to identify LRP1 ligands in human CSF that are differentially regulated by CR O-glycans. Our findings highlight the critical role of protein O-glycosylation in modulating functions of the LDLR/LRP endocytic receptors and demonstrate that the CR linker O-glycans tune the ligand preference of LRP1.

## Results

### O-glycans in CR linkers of LRP1

During the last decade we and others have created large datasets of human O-glycoproteomic data from cell lines and tissues, and these are available for perusal in the glyco.me database^23^. Here we surveyed this database to assemble data for O-glycans on human LRP1 that contain 27 CR linkers of which 10 conform to the **C_6_**X_3-5_**<u>T</u>C_1_** motif identified as the non-redundant sequence motif for GALNT11 mediated site-specific O-glycosylation^14,15^ (**Fig. 1a**). Our analysis revealed experimental evidence for O-glycans at 13 out of the 27 CR linker sequences in LRP1, with the highest frequency of O-glycans in the CR cluster II. Since our glycoproteomic data also include comparative analysis of wild-type and *GALNT11^-/-^* cells, we could confirm that eight of these O-glycosylated linkers, including the N-terminal linker-like motif preceding CR1, were only found O-glycosylated in the presence of GALNT11 activity^17,18^. This was further verified by data from our previous cell models with inducible expression of GALNT11, where gradient induction of GALNT11 tightly correlates with occupancy of O-glycans in LRP1 linker sequences^24^. Notably, the CR linker O-glycosylation sites are highly conserved throughout vertebrates (**Supplementary Fig. 1**) and also found in e.g. the orthologous lipophorin receptor in Drosophila^15^.

### O-glycosylation does not affect secretion of LRP1

To evaluate functions of GALNT11-mediated O-glycosylation of LRP1 we developed complementary human cell line models. The HEK cell line has low endogenous expression of LRP1, LRP2 and LDLR^24,16,25^ and was chosen as a model to introduce LRP1. We used two orthogonal methods to produce HEK cell lines with stable expression of full coding LRP1; (i) CRISPR/Cas9 mediated promotor <u>act</u>ivation of the endogenous LRP1 gene (Act; **Fig. 1b-c**) allowing for expression of LRP1 in its natural genomic context; and (ii) stable transfection of a <u>rec</u>ombinant cDNA construct encoding full-length LRP1 (Rec; **Fig. 1d**). We isolated stable clones (HEK-LRP1^Act#1^, HEK-LRP1^Rec#1^, HEK-LRP1^Rec#2^) by single-cell sorting and screening for LRP1 cell surface expression with anti-LRP1 antibodies selecting clones with similar expression levels (>10-fold higher surface staining compared to HEK WT) as evaluated by flow cytometry (**Fig. 1f**, **Supplementary Fig. 2**). Paired isogenic *GALNT11* knock-out (KO) clones (HEK-LRP1^Act#1-ΔT11^, HEK-LRP1^Rec#1-ΔT11^, HEK-LRP1^Rec#2-ΔT11^) were then produced by CRISPR/Cas9 KO of *GALNT11* (**Fig. 1. c, d**). Loss of GALNT11 activity did not affect LRP1 cell surface expression or subcellular localization (**Fig. 1e, f**), or the integrity of the LRP1 extracellular domain (**Fig. 1g**). We previously found that Lrp2 isolated from mouse kidney was recognized by Peanut agglutinin (PNA) lectin (binds core1 Galβ1-3GalNAc O-glycans without sialic acid) and that loss of Galnt11-mediated O-glycosylation substantially reduced binding^17^, which indicates that O-glycans of LRPs may be selectively undersialylated compared to other secreted glycoproteins that generally acquire O-glycans fully modified with sialic acids^26^. We therefore used SDS-PAGE Western blotting of cell lysates with lectins to evaluate effects of *GALNT11* KO on the synthesis of O-glycans on LRP1. We used the lectin Jacalin (binds unsialylated core1, core3 and Tn O-glycans^27^) to detect additional O-glycan structures. Probing with Jacalin demonstrated that LRP1 produced in cells with GALNT11 was strongly labelled, while LRP1 produced in cells without GALNT11 was only weakly labelled (**Fig. 1g),** which confirms the remarkable specific function of GALNT11 in directing the O-glycans on LRP1 and further suggests that these CR linker O-glycans are selectively undersialylated.

### GALNT11-mediated O-glycosylation modulates LRP1 ligand specificity and uptake

We previously showed that the loss of CR-linker O-glycans on LDLR and VLDLR reduces their binding to cognate lipoproteins, and similarly reduces Lrp2 binding to albumin^15,17^: LRP1 has a very broad ligand repertoire^9^, and we therefore developed high-throughput assays to quantitate cellular uptake of fluorescently labelled proteinaceous ligands using flow cytometry (**Supplementary Fig. 3a)**. We first monitored uptake of Alexa Fluor 488 labeled A2M, the canonical LRP1 ligand^28^, and 2N4R-tau (tau), a recently identified LRP1 ligand^22^, over a 2-hour period at 37°C in HEK-WT and HEK-LRP1 cells (**Supplementary Figs. 3, 4**). While A2M and tau was detectable in HEK-WT cells, LRP1 overexpression in HEK-LRP1 cells greatly increased the signal-to-noise ratio. For both HEK-LRP1^Act#1^ and HEK-LRP1^Rec#1/2^ a significant shift in median fluorescence intensity (MFI) could be measured within 30 minutes of ligand incubation and the signal increased linearly with time. At all timepoints ligand derived MFIs were approximately 10-fold (tau) or 20-fold (A2M) higher in HEK cells overexpressing LRP1 compared to HEK-WT cells, well in agreement with the 10-fold higher expression level of LRP1 compared to HEK-WT cells (**Fig. 1f**). Cells incubated with ligands for 2 hours at 4°C produced negligible signal compared to 37°C, indicating that the MFI changes detected at 37°C are derived from actively internalized proteins, which was confirmed by immunocytology (**Supplementary Fig. 5**).

To screen ligands for uptake by LRP1 expressing cells with and without GALNT11, we incubated cells with serially diluted ligands for 2 hours at 37°C. We included the addition of unlabeled RAP at 500 nM concentration for selective suppression of LRP-mediated endocytosis (**Fig. 2a)**. RAP is a LRP folding chaperone protein with low nanomolar binding affinity for LRP1^29^, and serves as a competitive receptor antagonist in this assay. Remarkably, uptake curves for two neurological ligands, tau and Aβ_1-42_, were differentially shifted in cells with KO of *GALNT11*, whereas the uptake of other similarly tested ligands (A2M, RAP, Apo-E2, -E3, -E4) was not affected (**Fig. 2a** and **Supplementary Fig. 6**). We measured tau uptake in a concentration span of 5-150 nM, and found that at all concentrations, the uptake of tau in HEK-LRP1^ΔT11^ cells was consistently reduced by 20% compared to the isogenic HEK-LRP1 cell clones with unaltered GALNT11 O-glycosylation capacity. The microtubule associated protein tau is a soluble phosphoprotein where hyperphosphorylation is associated with aggregation and neurological pathologies^30^. Interestingly, uptake in HEK-LRP1^ΔT11^ cells was further decreased to 25% of that of HEK-LRP1 cells when analyzing hyperphosphorylated tau (tau-Sf9), and 30% or 37% when analyzing phosphomimetic forms of tau containing either 3 or 9 lysine to glutamate substitutions, respectively (tau-3xKQ, tau-9xKQ) (**Fig. 2a**). In striking contrast to tau, uptake of Aβ in HEK-LRP1^ΔT11^ cells was increased by up to 30%. Testing a range of concentrations of fluorescently labeled monomeric Aβ_1-40_ and Aβ_1-42_, up to a maximum of 1,000 nM, we found that amyloid peptides were consistently taken up in a LRP1 and dose dependent manner with an increase in uptake of up to 30% for Aβ_1-42_ in all HEK-LRP1 cells with KO of *GALNT11* (**Fig. 2a**).

**Fig. 2.**
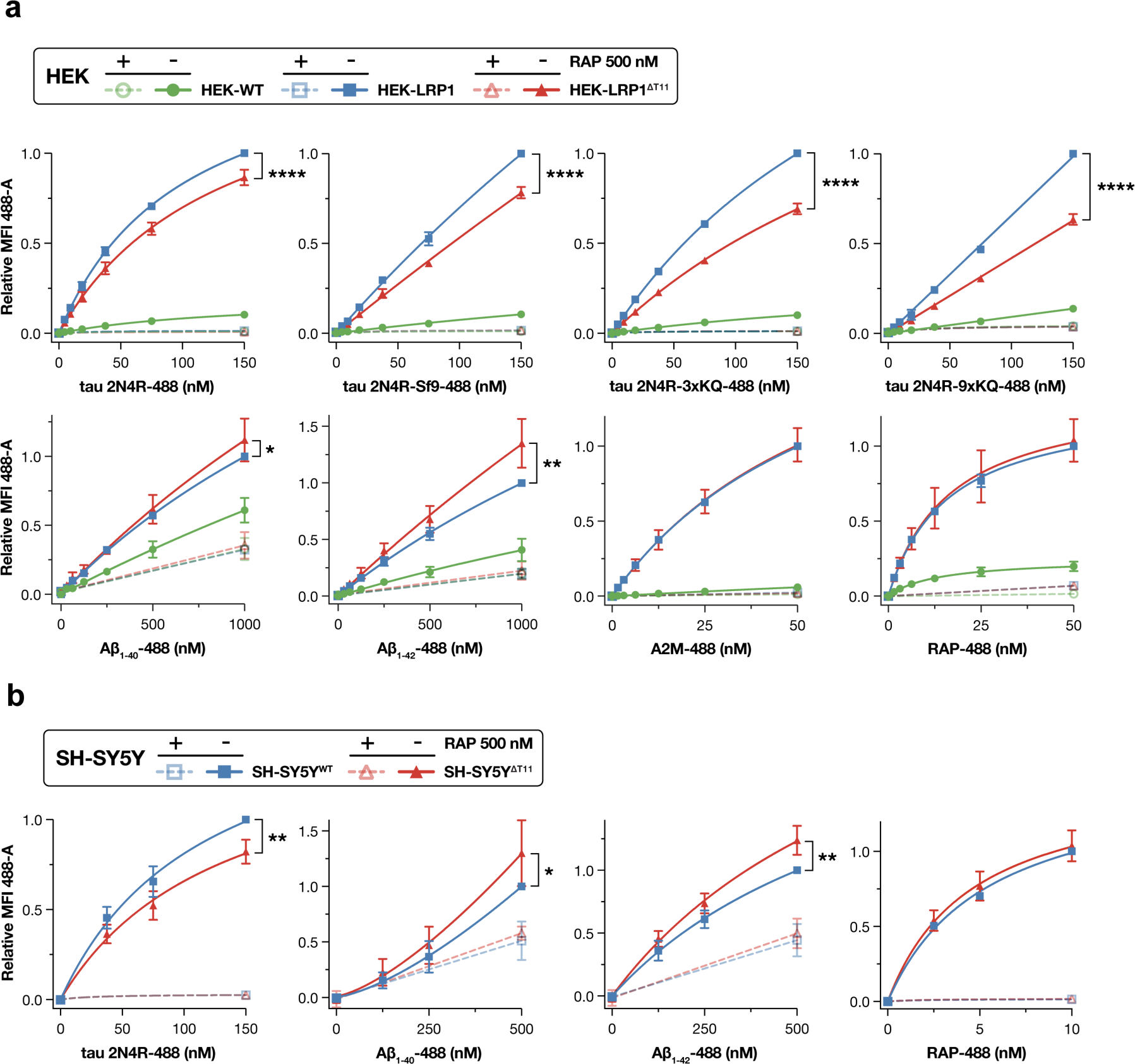
LRP1 ligand specificity and uptake is modulated by O-glycans initiated by GALNT11. **a** HEK-WT, HEK-LRP1 and HEK-LRP1^ΔT11^ cells were cultured overnight in a 96-well plate and the following day incubated with indicated Alexa or HiLyte Fluor 488-labeled ligands diluted in complete growth medium for 2 hours at 37°C. Cells were then washed, released and stained with propidium iodide (PI) before flow-cytometric analysis. The median 488-A fluorescence intensity (MFI) of the PI negative population was background subtracted using MFI values from matched cells cultured without ligand and thereafter normalized to the MFI recorded for HEK-LRP1 cells at the highest concentration of ligand used. Unlabeled RAP at 500 nM concentration was used to suppress LRP mediated endocytosis. Graphs represent the aggregate of three experiments, one for each of the HEK-LRP1^Act^, HEK-LRP1^Rec#1^ and HEK-LRP1^Rec#2^ cell lines. **b** Similar to above, SH-SY5Y^WT^ and SH-SY5Y^ΔT11^ cells were cultured in the presence of 488-labeled ligands for 2 hours at 37°C followed by flow-cytometric analysis, with the exception that for the Aβ_1-40_ and Aβ_1-42_ peptides these were applied to cells for 8 hours. Each graph represents the aggregate analysis of two experiments comparing SH-SY5Y^WT^ to three independent SH-SY5Y^ΔT11^ clones. Data are presented as mean ± SD and the ratio paired *t*-test was used assess significance (****P < 0.0001, ***P < 0.001, **P < 0.01, *P < 0.05).

We confirmed above findings in the SH-SY5Y neuroblastoma cell line using an isogenic pair of cells with and without GALNT11 expression. SH-SY5Y is a commonly used cell model to study uptake and degradation of tau and Aβ peptides^22,31–33^, and this cell line endogenously express LRP1 as well as GALNT11. We produced three independent *GALNT11* KO clones of SH-SY5Y (SH-SY5Y^ΔT11^) and applied the same protocol as above to measure uptake of tau, Aβ_1-40_, Aβ_1-42_ and RAP (**Fig. 2b**). In agreement with the findings in HEK cells, we found differential changes in uptake of tau and Aβ ligands. LRP1-dependent uptake of tau decreased by 20% in SH-SY5Y^ΔT11^, while the uptake of Aβ_1-42_ increased by 25% in SH-SY5Y^ΔT11^. We also confirmed that uptake of RAP remained unchanged.

### Discovery of LRP1 ligands in CSF affected by GALNT11-mediated O-glycosylation

The selective impact of O-glycosylation of LRP1 on ligand uptake prompted us to develop a method for unbiased discovery. LRP1 is expressed in choroid plexus epithelial cells lining ventricles where it is exposed to CSF, and LRP1 serves in selective uptake of CSF proteins destined for degradation or transcytosis across the CSF-blood barrier, including Aβ^34,35^. To identify LRP1-specific ligands affected by O-glycosylation, we labelled human CSF proteins with unnatural long-chain biotin (LC-biotin) for uptake in cells with or without GALNT11 and differential quantification by mass spectrometry (**Fig. 3a** and **Supplementary Fig. 7**). Across seven experiments utilizing three independent batches of pooled CSF, we identified 15 biotinylated proteins that were more abundant (P < 0.05) in cell lysates of HEK-LRP1^Act^ compared to HEK-WT cell lysates (**Fig. 3b** and **Supplementary Table 1**) and thus dependent on LRP1. None of these proteins exhibited differential uptake in the presence of RAP (**Supplementary Fig. 8**) and the three proteins CD14, MDH1 and ISLR represent potential novel LRP1 ligands. Furthermore, comparing HEK-LRP1^Act^ and HEK-LRP1^Act-ΔT11^ cell lysates, we identified seven proteins with different abundancies representing potential LRP1 ligands affected by O-glycosylation (**Fig. 3c** and **Supplementary Table 2**). The uptake of LC-biotinylated A2M displayed reduced uptake upon *GALNT11* KO (**Fig. 3c**) while fluorescently labeled A2M did not (**Fig. 2a**), and we hypothesize that the discrepancy in uptake observed could be due conformational and compositional differences of the two A2M preparations. The largest difference was observed for Hemopexin (HPX) where uptake was reduced by 40% in HEK-LRP1^Act-ΔT11^ cells compared to HEK-WT cells. We validated HPX as a ligand for LRP1 affected by GALNT11 dependent O-glycosylation using flow cytometry demonstrating that uptake of a fluorescently labeled HPX:Hemin complex was reduced by 20% in all three HEK-LRP1^Act/Rec#1/Rec#2-ΔT11^ lines (**Fig. 3d**).

**Fig. 3.**
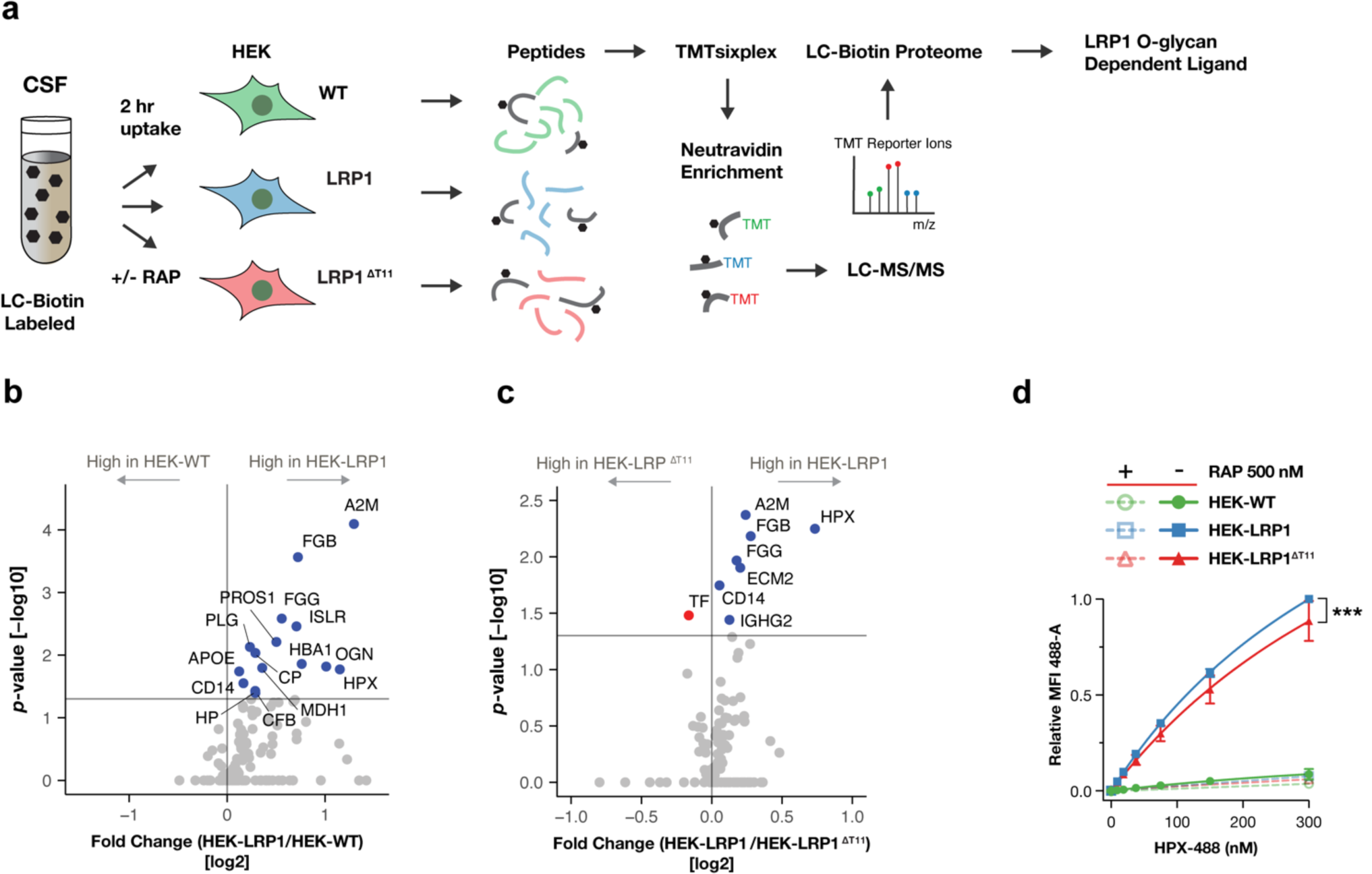
Cell-based discovery strategy for LRP1 ligands dependent on GALNT11-mediated O-glycosylation applied to human CSF. **a** Graphic representation of the workflow for the uptake assay. Cells with/without LRP1 overexpression and GALNT11-directed O-glycosylation capacity were cultured in DMEM supplemented with LC-biotin labeled human CSF proteins +/- 500 nM RAP at 37°C for two hours before being stringently washed and harvested. Following cell lysis, trypsin digestion, and TMTsixplex labeling allowing for direct comparison between cell lines, identical amounts of peptide samples were combined and subjected to neutravidin enrichment. After LC-MS/MS analysis, peptides modified with both LC-biotin and TMT were used to quantitate CSF protein uptake. **b-c** Volcano plots of CSF proteins identified across seven experiments utilizing three independent batches of pooled CSF with comparison of (**b**) HEK-LRP1^Act^ to HEK-WT and (**c**) HEK-LRP1^Act^to HEK-LRP1^Act-ΔT11^. Proteins significantly endocytosed (*P* < 0.05) are colored blue or red and annotated. **d** Flow cytometric measurement of Hemopexin:Heme complex uptake in HEK-LRP1 cells. Cells were incubated with Alexa Fluor 488-labeled Hemopexin:Heme diluted in complete growth medium for two hours at 37°C. The presented data represent the combined analysis of two experiments for each of the HEK-LRP1^Act^, HEK-LRP1^Rec#1^ and HEK-LRP1^Rec#2^ cell lines. Data are presented as mean ± SD and the ratio paired *t*-test was used assess significance (***P < 0.001).

### O-glycans on LRP1 are selectively hyposialylated

We previously reported that elongation and sialylation of the CR linker O-glycans attached by GALNT11 is required for enhanced affinity (>5-fold) of LDLR binding to LDL^15^. This finding places into question the observed apparent hyposialylation of Lrp2^17^ and LRP1 (**Fig. 1g**) and prompted us to in detail characterize the sialylation state of LRP1 O-glycans. Note that lectin enriched global O-glycoproteomics primarily reports *sites* of O-glycans in proteins, but rarely provide comprehensive information of the structures of O-glycans due to limitations of enrichment and analyses^36,37^. To circumvent these issues we therefore expressed and purified the soluble LRP1 ectodomain^38^ (Fig. 4a) to enable analysis of the mature GALNT11 directed O-glycans by mass spectrometry^39,40^. We first stably introduced a plasmid encoding the LRP1 ectodomain with C-terminal affinity purification tags into a HEK WT cell line with knock-in (KI) of RAP (HEK^KI^ ^RAP^) to generate HEK^KI^ ^RAP^-sLRP1 cells (**Fig. 4a, b**). We employed co-expression of RAP as this was demonstrated to improve secretion of soluble LRP1^38^. Like above, we then used CRISPR/Cas9 to KO *GALNT11* and generate isogenic HEK^KI^ ^RAP^-sLRP1^ΔT11^ cells. Comparable secretion of sLRP1 from HEK^KI^ ^RAP^-sLRP1 and HEK^KI^ ^RAP^-sLRP1^ΔT11^ cells was confirmed by Western blot analysis of conditioned media. Secreted sLRP1 and sLRP1^ΔT11^ proteins were purified from conditioned media by strep-affinity and size-exclusion chromatography (SEC), and analysis by SDS-PAGE revealed a dominant high molecular weight α-chain (>500 kDa) and the truncated β-chain (50 kDa). Lectin blot analysis confirmed that sLRP1 was highly labelled by Jacalin without prior removal of sialic acids by neuraminidase treatment (**Fig. 4c**), well in line with our results from labelling LRP1 in lysates from the HEL-LRP1^Act^ and HEK-LRP1^Rec^ lines (**Fig. 1g**). Mass-photometry and surface plasmon resonance (SPR) spectroscopy analysis of purified sLRP1 and sLRP1^ΔT11^ revealed a similar monomeric mass of around 600 kDa and both receptors bound RAP at low nanomolar concentrations in a 1:2 (sLRP1:RAP) binding stoichiometry, consistent with previous studies of transiently produced sLRP1^38^ (**Fig. 4d, e**).

**Fig. 4.**
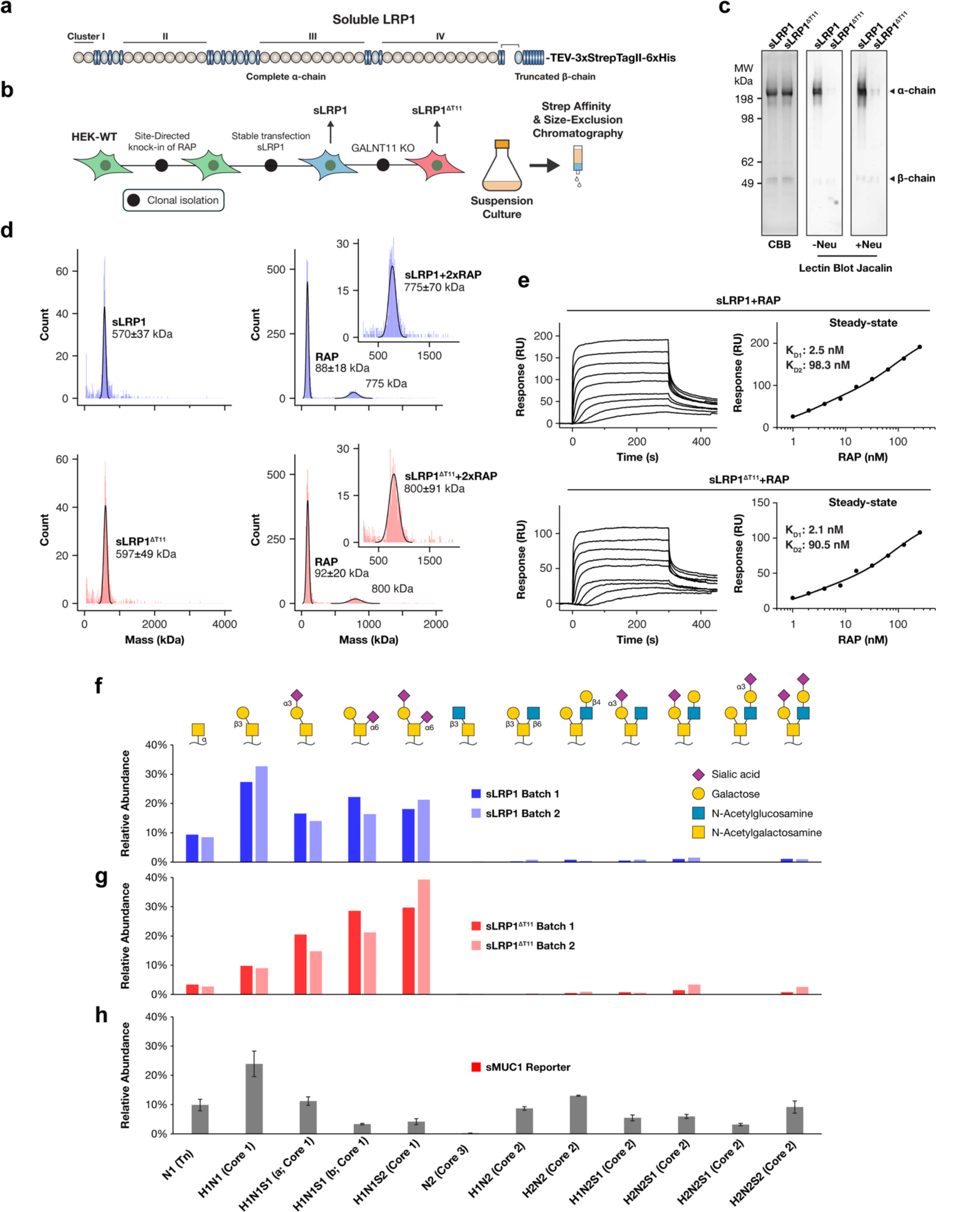
LRP1 CR O-glycans are hyposialylated. **a** Illustration of recombinant soluble LRP1 (sLRP1) protein where the transmembrane and intracellular domain of LRP1 have been replaced by a Tobacco Etch Virus (TEV) protease site followed by three tandem Strep-tag II sequences and a poly-histidine tag. **b** Lineage ancestry of clonal HEK cell lines engineered to stably express RAP and sLRP1 followed by knock-out of *GALNT11*. Cells were cultured in suspension and sLRP1 was purified from conditioned media by strep-affinity and size-exclusion chromatography. **c** Coomassie brilliant blue (CBB) and Jacalin lectin blot analysis of purified sLRP1 and sLRP1^ΔT11^ after non-reducing SDS-PAGE. Prior to incubation with Jacalin blotted membranes were treated with neuraminidase or buffer only. **d** Single molecule mass-photometry analysis of the complex formation between sLRP1 and RAP. **e** Surface plasmon resonance (SPR) analysis of sLRP1/sLRP1^ΔT11^ and RAP binding. The receptors were covalently immobilized to sensor surfaces and two-fold serial dilutions of RAP (1-256 nM) applied. Steady state K_D_ values were determined using a two-sites specific binding model. **f-h** LC-MS glycoprofiling of O-glycans released from two independent batches of purified (**f**) sLRP1, (**g**) sLRP1^ΔT11^ and (**h**) a single batch of a soluble MUC1 (sMUC1) reporter protein produced in HEK-WT cells^41^. Data are presented as the relative abundance of indicated O-GalNAc glycan. Three technical replicate analyses of sMUC1 were performed to demonstrate the variability of the assay and are presented as mean ± SD.

Finally, subjecting sLRPs to glycoprofiling, we found that the most abundant O-glycan on sLRP1 was non-sialylated core1 (H1N1, T; 30% relative abundance), followed by mono-sialylated core1 (H1N1S1-a and -b, mSTa and mSTb; 15% and 20%), di-sialylated core1 (H1N1S2, dST; 20%) and Tn (N1; 9%) (**Fig. 4f**). When analyzing sLRP1^ΔT11^ we observed a remarkable 3-fold decrease in the relative abundance of non-sialylated core1/T and Tn with a concomitant increase of mono- and di-sialylated core1/T (**Fig. 4g**). These results suggest that the O-glycans introduced in short CR linkers by GALNT11 are selectively prone for low sialylation, while O-glycans positioned at other sites, and introduced by other GALNTs, more faithfully follow the conventional sialylation efficiency for O-glycoproteins (**Supplementary Fig. 1**). O-glycans are generally found in unstructured regions of proteins such as linkers, but the short **C_6_**X_3-5_**<u>T</u>C_1_** linkers with the O-glycosite positioned -1 to the first disulphide-bonded cysteine residue (Cys1-Cys3) in-between the folded CR modules is likely to accommodate the unique substrate site for GALNT11 and also restrict further glycosylation efficiencies of the glycosyltransferases involved in elaboration of these O-glycans. The latter prediction is supported by our lack of detection of branched core2 O-glycans on sLRP1, which is widely found on other O-glycoproteins expressed in HEK cells, including a similarly produced soluble Mucin-1 (sMUC1) reporter^41^ (**Fig. 4h**).

### Sialylation of linker O-glycans are predicted to obstruct inter- and intramolecular interactions

The abundance of non-sialylated O-glycans in LRP1 CR linkers encouraged us to explore the structural and functional implications of sialylation of these linker O-glycans by molecular dynamics (MD) simulations. As a template we selected the NMR resolved structure of the non-glycosylated CR5-CR6 unit (that contains the **C_6_**X_3-5_**<u>T</u>C_1_** motif) in complex with RAP domain 1 (RAPd1; PDBid 2fyl) (**Fig. 5a**). The complex is formed by three networks of salt bridges involving the seven residue pairs between CR5-CR6 and RAPd1: Asp27-Lys24, Asp28-Lys24, Asp27-Lys60, and Asp28-Lys60; Asp66-Lys93, Asp68-Lys93 and Asp73-Arg34 (**Fig. 5b**). The non-glycosylated linker is disordered and extends away from the CR5-CR6:RAPd1 binding interface and into the solvent. We functionalized residue Thr44 in the C_6_AYP**<u>T</u>**C_1_ linker sequence with a core1 or monosialyl-core1 O-glycan structure (**Fig. 5c**) and ran a set of independent MD simulations. Three uncorrelated MD trajectories of 1 µs sampling time each revealed that while the glycosylated loop remains disordered, it can also fold over with the core1/T O-glycan forming stable interactions with Trp63 of CR6 and with the bound RAPd1. As shown in **Figure 5d**, the interaction hinges on a stacking CH-π interaction between the terminal Gal of the core1/T O-glycan and Trp63. Due to the weakness of this type of molecular interaction in additive, fixed-charge force fields, we probed its stability by augmenting the sampling, using this structure as a starting point and re-assigning random velocities. In this simulation the core1/T Gal residue shifts from the stacked conformation with Trp63 to engage in hydrogen bonding with RAPd1 Glu23 and Glu30 (**Fig. 5e**). Interestingly, these glycan dependent conformational changes could not be observed when the core1/T O-glycan was capped by a sialic acid residue (**Fig. 5c**). Two 1 µs MD simulations showed that the presence of the sialic acid residue does not allow stacking with Trp63 of LRP1, nor contact with RAPd1 due to steric hindrance. Furthermore, the two independent simulations lead to sampling of the same conformational space, where contact to the RAPd1 involves only CR5 and CR6, and where the linker remains disengaged.

**Fig. 5.**
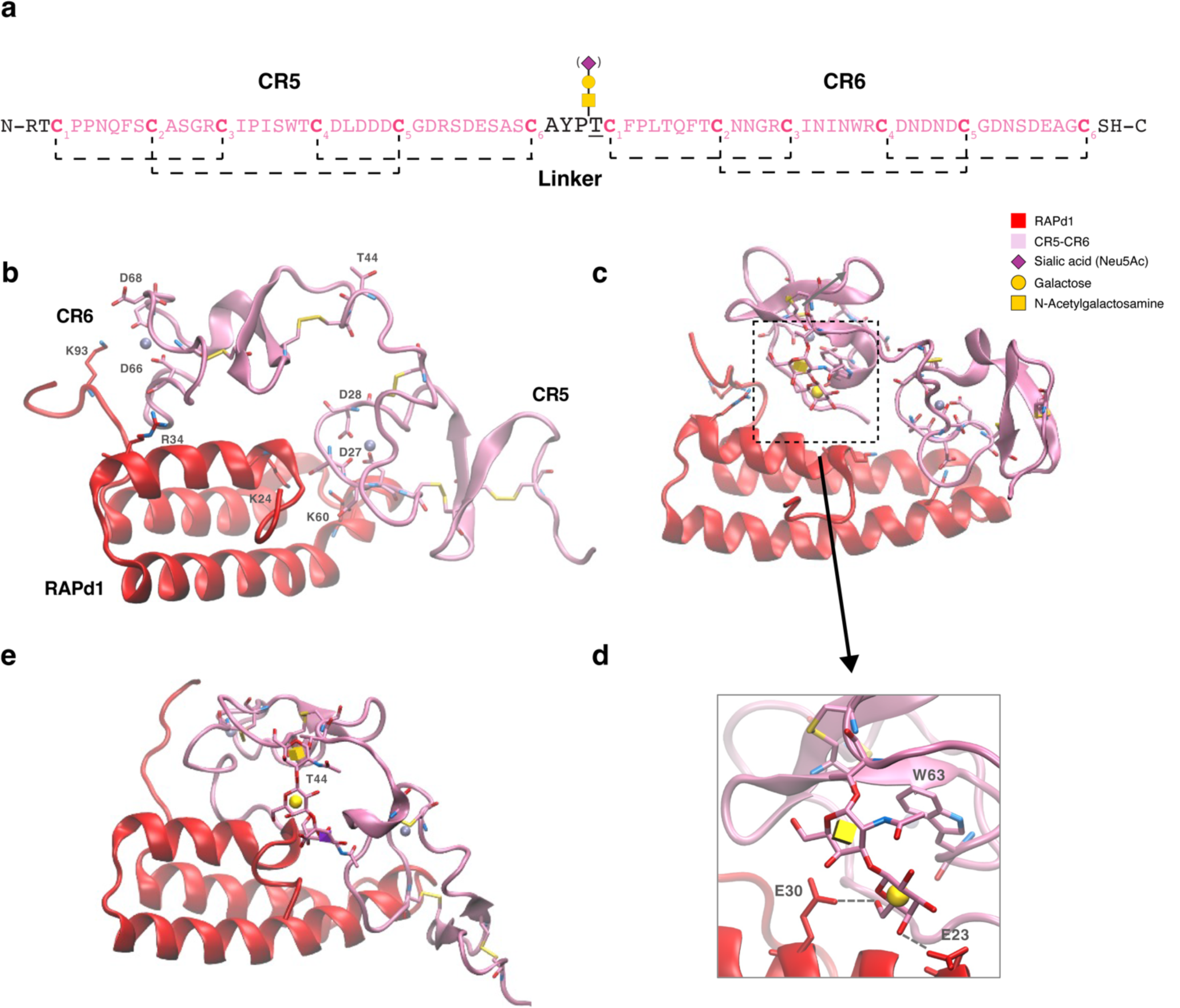
MD simulations of CR5-CR6 predicts the linker O-glycan modulation of inter- and intramolecular arrangements. **a** Primary sequence of the CR5-CR6 double domain with the short C_6_AYP**<u>T</u>**C_1_ linker sequence installed with a core1/T O-glycan with/without sialic acid capping. Intramolecular disulfide bonds are indicated by dashed lines. **b** NMR-guided docking structure of the complex between CR5-CR6 (pink) and RAP domain1 (red; PDB: 2fyl). **c** Representative structure from the MD simulations of the complex CR5-CR6:RAPd1 with an unsialylated core1 (T) O-glycan showing that the O-glycan interacts with Trp63 of CR6. **d** Expanded view of (**c**) displaying the hydrogen bond interactions between the core1/T O-glycan with the Glu23 and Glu30 residues of RAPd1. **e** Representative structure from the MD simulations of the complex CR5-CR6:RAPd1 with a sialylated core1 (mSTa) O-glycan showing that the O-glycan remains disengaged from the CRs and RAPd1 due to steric hindrance.

## Discussion

Here, we demonstrated that the ligand repertoire of LRP1 is modulated by O-glycans in the ligand-binding domains much like other LDLR/LRP endocytic receptors. We showed that CR O-glycans have distinct effects on LRP1’s binding and uptake of important neurological ligands including hemopexin, Aβ and tau. We confirmed that O-glycosylation of LRP1 is regulated by the GALNT11 isoenzyme and provided evidence that the linker O-glycans are selectively hyposialylated in cells. We previously demonstrated tight dose-dependent GALNT11 enzyme regulation of O-glycan occupancy on LDLR/LRPs, and here provide compelling MD simulation data to suggest that linker O-glycans play a key role in regulating conformations of the ligand-binding CR modules and hence the binding properties of LRP1. Finally, we generated and validated a cell-based platform for unbiased discovery of LRP1 ligands that are selectively regulated by CR linker O-glycans. LRP1 is widely expressed with a plethora of tissue-specific endocytic functions and a large and diverse repertoire of ligands, but how LRP1 serves so many ligands and biological functions remains unclear^9^. LRP1 contains four ligand-binding regions comprised of CR clusters I-IV with varying number of CRs (2-10 CRs), and while all CR clusters contain one or more linkers with identified O-glycans, linkers in cluster II stand out with near complete modification by O-glycans (**Fig. 1a**). CR clusters II and IV appear to contribute the most to ligand binding^9,42^. Studies with LRP1 mini-receptors encoding individual clusters have identified a broad range of ligands targeting this cluster II, including tau and Aβ^9,22^. LRP1 was recently identified as a master regulator of tau uptake and spread in the brain^22^, and here we found that LRP1-mediated uptake of tau was significantly lower in the absence of CR linker O-glycans (20%; **Fig. 2**). LRP1 also participates in clearance of Aβ and in regulating brain Aβ metabolism in neurodegenerative disorders^9,43^, and we found that loss of CR linker O-glycans significantly improved Aβ uptake (**Fig. 2**). GALNT11 is similar to LRP1 expressed in most cells and tissues, and thus GALNT11 activities provides a compelling regulatory mechanism for fine-tuning the ligand-binding properties and ligand repertoire of LRP1 as have been described for the tissue-specific LDLR^15^ and LRP2 receptors^17,24,44,45^.

The aggregation of extracellular Aβ and deposition of intracellular hyperphosphorylated tau are the pathological hallmarks of Alzheimer’s disease (AD)^46^. The initiation of AD is attributed to a sustained imbalance between Aβ production and clearance, leading to the accumulation of toxic and inflammatory Aβ plaques, which in turn trigger neuronal production of hyperphosphorylated tau capable of seeding and spreading tau tangles^30^. While LRP1-mediated clearance of extracellular Aβ is generally viewed as protective against AD, the role of LRP1 clearance in tauopathies is ambiguous^22,31^. Cooper and colleagues proposed that LRP1 is important for the removal and degradation of normal tau as well as the spread and seeding of pathological and posttranslationally modified tau variants^31^. This study also reported LRP1 binding affinities to tau and found that both hyperphosphorylation and phosphomimetic lysine to glutamate substitutions reduced affinity, establishing the binding hierarchy: tau > tau-sf9 > tau-3xKQ > tau-9xKQ^31^. Interestingly, we find that the glycosylation dependence of tau variants is inversely correlated to their LRP1 binding affinity (**Fig. 2a**), where weaker binding variants like phosphorylated tau are more sensitive to the loss of CR linker O-glycans. Collectively our data suggests that GALNT11-directed O-glycans contribute to LRP1 uptake of tau, in particular hyperphosphorylated variants of tau, while O-glycans reduce the uptake of Aβ peptides. This suggests that O-glycans have the potential to contribute to the progression of AD by differentially influencing LRP1 interaction with these two ligands.

While O-glycans in CR linkers are critical for the endocytic functions of LDLR/LRPs, further studies will determine how O-glycans mediate these effects. Based on our previous finding, O-glycan elongation, beyond the initial GalNAc (Tn), is required to produce LDLR variants with a high affinity to LDL^15^. Our findings here with LRP1, and previously with Lrp2^17^ identifies a Core1 extended structure, with a generally incomplete sialic acid capping, as an abundant linker O-glycan. In contrast, core1 O-glycans with complete sialic capping are the predominant structures found on most glycoproteins produced in cells lines, including HEK, CHO and SH-SY5Y^25^ (Madsen et al in review), which suggests that the incomplete sialic acid capping on LDLR/LRP linker O-glycans is atypical and likely related to the specific O-glycosylation pathway regulated by GALNT11 and/or the specific topological context of these O-glycans being positioned in short linkers in-between the folded CRs. It is also possible that interactions between LDLR/LRPs and their obligate chaperone RAP affects the O-glycosylation and in particular sialylation during Golgi trafficking. RAP is proposed to dissociate from nascent LRP1 in medial Golgi compartments in a pH dependent manner prior to LRP1 transport to the cell surface^47^. Our MD simulations of LRP1 CR5-CR6 in complex with RAP suggest that O-glycan modification with sialic acid do affect the conformation of CRs by abrogating interactions between the core1 O-glycan and the C-terminal CR. Specifically, we found that without an O-glycan the linker is solvent exposed and that the installation of a core1 (T) O-glycan (by the GALNT11 and C1GALT1 glycosyltransferases) provides a mechanism by which the linker can engage the LRP1:RAP interface thereby possibly preventing further extension or sialylation of the glycan as long as RAP remains bound. Interestingly, simulations with a sialylated core1 O-glycan showed that these interactions are incompatible with sialylation due to steric hindrance. The main sialyltransferase isoenzymes involved in capping core1 O-glycans are ST3GAL1 and ST3GAL2, where ST3GAL1 is suggested to be active in both medial and trans-Golgi compartments^29,48^, and whether LRP1 linker O-glycans remain inaccessible to ST3GAL1 after RAP dissociation due to LRP1 structural features or differential compartmentalization of LRP1 and ST3GAL1 is not known. Further studies are needed to elucidate the timings and possible overlaps of RAP and GALNT11 functions during the biogenesis of LRP receptors.

It is important to note that the sialylation state of membrane glycoproteins is at least partly subject to dynamic regulation. Membrane proteins have long been known to undergo some degree of sialylation (or resialylation) during constitutive recycling^49^. More recently a novel growth factor induced mechanism referred to as a GlycoSwitch was proposed to rapidly mediate endocytosis and retrograde trafficking of membrane glycoproteins to the Golgi and (re)exposure to the glycosylation machinery^50^. Interestingly, this GlycoSwitch functions through a local pH drop at the cell surface and activation of cell membrane resident neuraminidases (NEU1/3) to trigger galectin-mediated endocytosis of select glycoprotein clients. LDLR/LRPs are possibly clients of this GlycoSwitch^50^, which could provide for rapid and dynamic changes in O-glycan occupancy, structures, and their sialylation state through growth factor regulation. In support of this hypothesis is the finding that the apparent hyposialylation of Lrp2/megalin we find in the normal mouse kidney^17^, was indeed eliminated in mice deficient in the *Neu1* gene^51^, suggesting that NEU1 plays a role in regulating the sialylation state of O-glycans on LRP2 and other receptors. How this occurs warrants further studies

Our analysis of glycoproteomic data revealed that most **C_6_**X_3-5_**<u>T</u>C_1_** consensus sequence CR linkers in LRP1 have O-glycans attached (**Supplementary Fig. 1**), and in a recent cryo-electron microscopy (cryo-EM) study of LRP2 O-glycans were found in 70% of the linkers^52^. Notably, both these methodologies (lectin enriched O-glycoproteomics and cryo-EM) can be used to identify positions of O-glycans but not the specific O-glycan structure attached. The former method involves lectin-based enrichment of specific O-glycan structures^36^, and as glycans are flexible Cryo-EM can only assign the inner monosaccharides^52^. However, as mentioned above, studies using lectin binding clearly point to selective incomplete sialic acid capping of LRPs in cells and tissues. In lines with this we previously found that PNA reacted strongly with Lrp2 isolated from wildtype mouse kidneys, and PNA reactivity on mouse kidney tissue sections co-localized with Lrp2^17^. Moreover, this PNA reactivity was independent of tissue pretreatment with neuraminidase to remove sialic acids^17^, and absent in kidneys from *Galnt11*^-/-^ mice^17^. Here, we extended these findings to the characterization of the complete extracellular domain of LRP1 produced by HEK cells. Using Jacalin lectin labelling and released O-glycan profiling, we confirmed that a majority (90%) of O-glycans are core1 structures and a that large fraction of these are not modified with sialic acid^27^ (**Fig. 1g, 4b, f, g**). Our LRP1 ligand uptake studies were performed in HEK cells with their steady-state incomplete sialylation capacity, and it is conceivable that the effects observed on the ligand repertoire will be exacerbated with no or more complete sialylation of LRP1. Further studies of regulation of O-glycosylation and sialylation of the CR linkers in all LDLR/LRP endocytic receptors are clearly needed, but we predict that regulation of these O-glycans perhaps through the novel GlycoSwitch provides for rapid and dynamic fine-tuning of the affinities and ligand repertoire of these important receptors to meet changing physiological requirements.

We previously reported progressive reduction of Lrp2 protein in kidney proximal tubule cells in aging *Galnt11*^-/-^ mice, which ultimately resulted in kidney decline^17^. More recently, associated effects from loss of LRP kidney function on vitamin D homeostasis in this mouse model was described^53^. These results are in line with studies identifying GALNT11 as a GWAS candidate for chronic kidney decline^54^. In our cell model studies of expression of LDLR^15^, Lrp2^17^, and here LRP1, we have not observed aberrancies in secretion or cell surface expression/stability of these receptors in cells with KO of *GALNT11*. While loss of Lrp2 in proximal tubule cells in *Galnt11* deficient mice may have several causes, we find in vivo receptor exhaustion and/or protein toxicity associated with impaired Lrp2 functions due to loss of O-glycans to be among the most compelling reasons. The systemic loss of Galnt11 in this mouse model did result in loss of CR linker O-glycosylation of LDLR/LRP receptors in all tissues analyzed, and indeed we observed increased plasma LDL-cholesterol in *Galnt11^-/-^* mice compared to wild-type suggesting dysfunction of Ldlr^17^. More detailed analysis of effects on distinct LDLR/LRP receptors should be studied in a tissue specific context in conditional-null mice. Moreover, given our findings here that LRP1-dependent Aβ uptake is decreased by O-glycosylation of GALNT11 (**Fig. 2**) it is noteworthy that LRP1-mediated transcytosis is a major pathway for removal of Aβ peptides from brain^55–59^. Unlike tau, we could not fully suppress Aβ peptide internalization by co-incubation with RAP, likely due to alternative cellular entry routes for these peptides^21^. However, the increased Aβ uptake in both HEK-LRP1^ΔT11^ and SH-5YSY^ΔT11^ was antagonized by RAP and we therefore hypothesize that GALNT11-mediated O-glycans on LRP1 are important for Aβ uptake and transcytosis at the blood-brain and blood-CSF barriers^55^.

The ligand repertoire of LRP1 has been studied in great detail^9,60^, and here we developed an isogenic HEK cell system and workflow for differential discovery of LRP1 ligands that are affected by GALNT11 O-glycosylation and applied this to proteins of CSF (**Fig. 3a**). LRP1 is expressed in several cell types in the brain, including in the choroid plexus epithelium (CPE) responsible for maintaining CSF homeostasis. Although we do not fully understand the roles of LRP1 in the CPE, multiple studies have demonstrated that LRP1 clear Aβ from CSF and deliver Aβ to the blood by transcytosis, and it is likely that LRP1 serves other ligands similarly, much like the role of LRP2 in the kidney proximal tubule epithelium^12^. Here our mass spectrometry-based strategy for assessing the role of LRP1 O-glycans in interactions with human CSF proteins resulted in the identification of several LRP1 ligands, both known and novel, with and without sensitivity for GALNT11 introduced O-glycans. We identified Hemopexin as depending on O-glycans for uptake. Hemopexin functions to capture free heme released during hemoglobin breakdown after which the complex is internalized via LRP1 thus protecting cells and proteins from heme induced oxidative damage and maintaining iron homeostasis^61^. Heme is implicated in AD and can directly bind to Aβ to form a complex capable of neurotransmitter oxidation as well as altering Aβ aggregation properties^62,63^, and low levels of CSF hemopexin has been associated with early stage AD^64^.

In conclusion, our study demonstrates that the endocytic functions of LRP1 are regulated by O-glycosylation and provides compelling evidence that this regulation affects the metabolism and turnover of hemopexin, tau and Aβ. These findings position O-glycosylation of LRP1 at the nexus of the metabolic and clearance defects associated with neurodegenerative diseases and provides impetus for further studies in e.g. conditional mouse models. Thus, O-glycosylation initiated by GALNT11 introduces a new level of regulation for the complex LDLR/LRP receptors that may serve in rapid responses to physiological stimuli through remodeling of existing cell surface located receptors.

## Methods

### Cell culture

HEK293-6E (HEK; obtained through a license agreement with Dr. Yves Durocher, Bioprocédés Institute de recherche en Biotechnologie, Montréal) and clones produced herein were cultured in DMEM supplemented with 10% fetal bovine serum (FBS) and 2 mM GlutaMAX (Gibco™, Thermo Fisher) in a humidified incubator at 37°C and 5% CO_2_. The Neuroblastoma Cell Line SH-SY5Y and all isogenic clones were cultured in a 1:1 mixture of RPMI 1640 and DMEM supplemented with 10% FBS and 2 mM Glutamax at 37°C and 5% CO_2_.

### Plasmids for recombinant expression of LRP1

LRP1 (Uniprot Q07954 with the following mutations; N603I and Q2900P) cDNA without signal peptide or stop codon and cloned into pCR8/GW/TOPO (Invitrogen™, Thermo Fisher) was kindly provided by Dr. Piet Gros^38^. We modified this plasmid by annealed oligo cloning to introduce the native LRP1 signal peptide and a C-terminal *2xFLAG-PacI-STOP*. The resulting LRP1-2xFLAG open reading frame was subsequently *NotI* subcloned into *pIRES-Puro3* (Takara) generating *pIRESpuro3-LRP1-2xFLAG* encoding full-length LRP1(aa1-19, ARS, aa20-4544)-2xFLAG. To generate a plasmid expressing a soluble form of LRP1 we *SgrAI/PacI* excised a fragment encoding the transmembrane domain and downstream parts of LRP1 from *pIRESpuro3-LRP1-2xFLAG* and replaced it with a dsDNA fragment (Genewiz, USA) encoding a TEV protease cleavage site and affinity purification tags to produce the plasmid *pIRESpuro3-sLRP1-TEV-3xStrepTagII-6xHis* that encodes LRP1(aa1-19,ARS, aa20-4409). All plasmid sequences were verified by Sanger sequencing.

### Stable activation of *LRP1* by EF-1ɑ promoter knock-in

A gRNA (cccctggtgcgctttgccga) binding 140 bp upstream of LRP1 start codon was identified using the ChopChop tool^65^ and its activity validated by Indel Detection by Amplicon Analysis (IDAA) as described previously^66^. A PCR fragment encoding human EF-1ɑ promoter was amplified from an overexpression plasmid using CloneAmp HiFi PCR Premix (Takara 639298) and gel purified. HEK-WT cells were seeded in 6-well plates and the next day transfected with 1 µg of PBKS-Cas9-2A-eGFP plasmid (Addgene #68371), 1 µg gRNA plasmid, 1 µg EF-1ɑ promoter PCR fragment and 1 µg of NOE scrambled oligo using Lipofectamine 3000. The day after transfection, EGFP positive cells were bulk sorted by FACS (SONY SH800). After expanding the bulk-sorted population for two weeks cells were rinsed with PBS, resuspended by pipetting in PBS 0.5% serum albumin (BSA), stained with anti-LRP1 mAb 8G1 followed by goat anti-mouse Alexa Fluor 647 and single cell sorted, gating for EGFP negative and LRP1 positive cells (top 1%). Sorted clones were expanded and screened by PCRs designed to span the junctions of LRP1 3’UTR and the EF-1ɑ promoter. Antibodies, lectins and streptavidin conjugates used in this study were diluted according to Supplementary Table 3.

### Knock-in of RAP in HEK cells

Site-directed KI of CMV-RAP at the AAVS1 safe harbor locus was performed using a modified ObLiGaRe gene KI strategy as previously published^24,67^. An EPB71 knock-in donor plasmid containing a cassette encoding CMV driven RAP followed by a BGH polyA terminator flanked by inverted AAVS1 Safe-Harbor locus ZFN binding sites was produced by In-Fusion (Takara) subcloning of RAP cDNA from pQTEV-LRPAP1 (Addgene 31327) into EPB71. KI clones were screened by PCR with primers designed to span the junction of the donor casette and the AAVS1 locus. A mono-allelic KI clone was selected for stable transfection of *pIRESpuro-sLRP1-3xTEV-StrepTagII-6xHis*.

### Generation of HEK clones expressing recombinant full-length or soluble LRP1

HEK-WT cells and a HEK^KI^ ^RAP^ clone were seeded in 24-well plates, and the following day transfected with 250 ng of *pIRESpuro-LRP1-FLAG_2_* and *pIRESpuro-sLRP1-StrepTagII_3_-His_6_*, respectively, using Lipofectamine 3000 according to the manufacturer’s instructions. The following ten days cells were put under selective pressure in medium supplemented with 1 µg/mL of puromycin and thereafter maintained in the presence of 0.25 µg/mL puromycin. Bulk-selected cells were single-cell sorted by FACS (Sony SH800). Clones stably transfected with *pIRESpuro3-LRP1-2xFLAG* were screened for LRP1 expression by immunocytology microscopy using anti-LRP1 mAb 8G1 and anti-Mouse IgG Alexa Fluor 488 secondary antibody. Supernatants from clones transfected with *pIRESpuro3-sLRP1-3xStrepTagII-6xHis* were screened by LRP1 western blot as described below.

### CRISPR/Cas9-targeted KO of *GALNT11*

Cells cultured in 6-well plates to ∼70% confluency were transfected with 1 µg PBKS-Cas9-2A-EGFP plasmid and 1 µg *GALNT11* gRNA plasmid^68^ using Lipofectamine 3000. The day after transfection cells were bulk sorted by FACS (SONY SH800) based on EGFP expression. After one week in culture, bulk-sorted cells were single-cell sorted into 96-well plates. Expanding clones were screened by IDAA to identify frameshift mutations and candidate KO clones were validated by sanger sequencing (Supplementary Table 4).

### LRP1 western and lectin blot

Cells cultured in 24-well plates were rinsed with cold PBS and lysed on ice in modified RIPA buffer (50 mM Tris pH 8, 150 mM NaCl, 1% Triton X-100, 0.5% Na Deoxycholate, 1 mM EDTA) supplemented with cOmplete™ Protease Inhibitor Cocktail (Roche), for 20 min under gentle agitation. Lysates were cleared by centrifugation (12,000 g, 10 min, 4°C) and protein concentration determined by BCA assay (Thermo Fisher 23225). To blot the LRP1 α-chain clarified total cell lysates or samples containing recombinant soluble LRP1 were combined with 2x Tris Glycine sample buffer without reducing agent and heated for 10 min at 82°C. Samples were separated on 4-12% Novex Tris-Glycine gels using Tris Glycine SDS running buffer (Invitrogen™ Novex™ system) and transferred to PVDF membranes utilizing the iBlot2 (Thermo Scientific) dry-blot transfer device for 11 minutes at 20 V. For Jacalin lectin blot, membranes were blocked in a 1% solution of Polyvinylpyrrolidone (40,000 MW) in Tris Buffered Saline with 0.05% Tween 20 (TBS-T) at 4°C overnight followed by incubation with neuraminidase (Sigma N2876) diluted in PBS for 1 hour at 37°C prior to 1-hour incubations at room temperature with biotinylated Jacalin and Streptavidin HRP diluted in TBS-T. For LRP1 ɑ-chain western blot, membranes were blocked in a 5% solution of skim-milk in TBS-T, followed by incubation with mouse anti-LRP1 mAb 8G1 and rabbit anti-mouse HRP conjugated secondary antibody, both diluted in TBS-T. Blots were developed using SuperSignal™ West Pico PLUS Chemiluminescent Substrate (Thermo Fisher) and imaged on a ImageQuant™ LAS 4000 (GE Healthcare).

### Immunocytology

25,000 cells were seeded on glass coverslips placed in 24-well plates and the following day cells were fixed in 4% paraformaldehyde for 15 minutes, rinsed with PBS and permeabilized in block buffer (PBS with 5% FBS, 0.5% Triton X-100, and 50 mm glycine) for 30 min at room temperature. For non-permeabilized conditions Triton X-100 was omitted from the block buffer. Primary and secondary antibodies were diluted in block buffer and sequentially incubated for 1 h at room temperature with PBS washes. After the final PBS wash coverslips were mounted with ProLong™ Gold Antifade supplemented with DAPI (Thermo Fisher P36931) and imaged on an Axio Imager upright fluorescence microscope (Zeiss) fitted with an Axiocam 503 camera (Zeiss), and a 40x/0.8 NA Plan-Apochromat and Plan-Apochromat 63x/1.40 Oil objective. Image acquisition adjustments was performed using ZEN software (Zeiss).

### Flow-cytometric measurement of LRP1 surface expression

Cells cultured in 24-well plates were rinsed with PBS and released by pipetting into PBS, 0.5% BSA. Cell suspensions were transferred into wells of V-bottom 96-well plates and centrifuged (350g, 3 minutes, 4°C) and washed once by resuspension in PBS, 0.5% BSA. Cells were incubated with anti-LRP1 mAb 8G1 diluted in PBS, 0.5% BSA for 1 hour at 4°C, washed twice, and incubated with anti-Mouse IgG Alexa Fluor 488 diluted in PBS, 0.5% BSA for 1 hour at 4°C followed by two washes. Cells were run on a SONY SA3800 flow-cytometer and FCS files analyzed using FlowJo (v10.10).

### Purification and labeling of proteins used in LRP1 uptake assays

Full length tau protein (2N4R, 1-441aa) was expressed in *E. coli* Rosetta 2 transformed with *pET29b-Tau* (Addgene 16316) and purified with slight modification to previously published protocols^32,69^. Cell pellets were resuspended in cell lysis buffer (50 mM MES pH 6.5, 5 mM DTT, 1 mM PMSF, 1 mM EGTA) + cOmplete protease inhibitor tablets (Roche), boiled for 20 minutes, and then centrifuged at 13,000 × g for 20 min at 4°C. The supernatant was filtered and loaded onto a MonoS column (Cytiva) equilibrated with Buffer A (50 mM MES pH 6.5, 50 mM NaCl, 2 mM DTT, 1 mM PMSF, 1 mM EGTA) and eluted with a linear gradient of Buffer B (Buffer A + 1 M NaCl). Fractions containing 2N4R tau were pooled and polished by size exclusion chromatography on a Superose 6 Increase equilibrated in PBS. His-tagged tau variants 2N4R-3xKQ and 2N4R-9xKQ, his-tagged phosphorylated 2N4R tau and his-tagged Receptor-associated protein (RAP) were produced as described previously^31,70^.

The pET32-E43C vector encoding human ApoE4 fused to Thioredoxin (Trx) was kindly provided by A. Chroni (National Center for Scientific Research, Greece), and used as a template for the generation of ApoE2 and ApoE3 encoding vectors by site-directed mutagenesis (GenScript). ApoE isoforms were expressed in *E. coli* BL21 (DE3) pLysS and purified as described previously^71^. Briefly, ApoE-Trx was isolated from clarified cell lysates by nickel-affinity chromatography and separation of Trx from ApoE was performed in 20 mM Tris-HCl, pH 8.0, 0.3 M NaCl and 1 mM DTT using 3C-protease with a 3C-protease to Trx-ApoE ratio of 1:25 (w/w) for 18 h at 4°C(). Liberated ApoE was isolated by nickel-affinity chromatography performed in same buffer with the addition of 6M urea to aid in complex dissociation. ApoE was recovered from the column flow-through and dialyzed against 5 mM NH_4_HCO_3_ before being lyophilized and stored at -80°C until use.

Alpha 2 Macroglobulin fast form from human plasma was acquired from Sigma (SRP6315). Hemopexin from human plasma and hemin was acquired from Sigma (H9291-500UG and 51280-1G)and combined at a a two-fold molar excess of Hemin for 2 hours at 37°C before being desalted using a ZebaSpin column equilibrated in PBS. All proteins were labeled with Alexa Fluor 488 NHS ester (Thermo Fisher A20000) according to the manufacturer’s instructions. Protein concentration was determined using BCA assay (Thermo Fisher 23225). Aβ_1-40_ and Aβ_1-42_ tagged with an N-terminal HiLyte Fluor™ 488 moiety were ordered from Anaspec (AS-60491-01, AS-60479-01) and resuspended in 1,1,1,3,3,3-Hexafluoro-2-propanol (HIFIP), aliquoted, dried, and stored in a desiccator at -20°C. Before use in cellular uptake assays peptides were solubilized in DMSO by water bath sonication.

### Flow-cytometric measurement of LRP1 ligand uptake

HEK cells were seeded at 20,000 cells per well and SH-SY5Y cells were seeded at 50,000 cells per well in polyethyleneimine (Polysciences PEI25K) coated flat bottom 96-well plates. The next day the medium was replaced, and cells were treated with 488-labeled proteins diluted in complete growth medium or Opti-MEM (Gibco) for 2 h at 37°C (unless otherwise indicated in the figure legends). Cells were then rinsed with PBS, trypsinized and resuspended in complete growth medium, and transferred to a round bottom 96-well plate placed on ice. Propidium iodide (PI) was added to wells at a final concentration of 1 ug/mL and cells were immediately analyzed on a SA3800 Flow Cytometer (SONY) equipped with a chilled plate loader. HEK-WT, HEK-LRP1 (Act#1, Rec#1 and Rec#2) and corresponding HEK-LRP1^ΔT11^ clones were run on the same plate and at least 5,000 singlet and PI negative cells were recorded from each well. Control wells with untreated cells were used to establish the background 488-A MFI for each cell line. To calculate relative MFI 488-A, background subtracted values were normalized to HEK-LRP1 or SH-SY5Y^WT^ uptake for the highest ligand concentration used. The clone HEK-LRP1^Rec#1-ΔT11^ exhibited a 20% increase in LRP1 surface expression compared to HEK-LRP1^pCMV#1^. Therefore, all relative MFI 488-A values from LRP1^Rec#^ ^1-ΔT11^ were adjusted by a factor of 1.2. Automated gating and analysis was performed in R using the packages openCyto^72^ and flowCore^73^. To block LRP uptake, we included unlabeled RAP at 500 nM concentration. To control for cell surface binding, duplicate plates were prepared and briefly placed on ice before the addition of chilled ligand containing medium and incubation at 4°C.

### LC-MS/MS based identification and quantification of endocytosed CSF proteins

CSF was collected from patients admitted to the Trauma Centre at Copenhagen University Hospital on suspicion of a subarachnoid hemorrhage and stored at -80°C. Samples from patients with negative findings, including a negative CT scan taken 6 hours after symptoms, and where no blood was found in the CSF were used for experiments. All samples were anonymized and pooled to ensure no patient traceability. To 9 volumes of pooled CSF was added with 1 volume of 200 mM HEPES buffer pH 7.4 and 0.1 volumes of 320 uM NHS-LC-Biotin (Thermo Fisher 21336) dissolved in DMSO. The mixture was incubated rotating end over end for 2 hours at room temperature, buffer exchanged into HEPES buffered DMEM medium using Zeba Spin desalting columns (Thermo Fisher), aliquoted and snap frozen in an ethanol dry ice bath. For uptake assays, 3.5 million HEK cells were seeded 10 cm dishes coated with PEI and the following day the medium was replaced to incubate cells with 3.5 mL of biotinylated CSF in DMEM for 2 hours at 37°C. Dishes were then moved to a cold room and rinsed two times with chilled PBS and two times with chilled 0.1 M glycine pH 3, 150 mM NaCl. The cells were then released into PBS by scraping, collected by centrifugation at 500×g for 5 min at 4°C and stored at -80°C. To block LRP uptake we included dishes where the biotinylated CSF in DMEM was supplemented with 500 nM RAP. To control for cell surface binding, we included dishes that were briefly placed on ice before the addition of chilled biotinylated CSF in DMEM and incubation at 4°C. Total cell lysates were digested with trypsin and peptides labeled with tandem mass tags (TMTsixplex, Thermo Fisher) as described previously^24,74^. From each of the six samples 0.2 mg of labeled peptides were combined, dried in a speed-vac, solubilized in 1 mL of PBS and incubated with 25 µl of settled Pierce™ NeutrAvidin™ UltraLink™ Resin (Thermo Fisher 53150) for 1 hour at room temperature in a micro spin-column. The resin was washed six times with 0.5 mL PBS, 5% Acetonitrile and once with 0.5 mL ultrapure water followed by five elution steps with 0.5 mL 80% Acetonitrile, 0.2% trifluoroacetic acid (TFA), 0.1% formic acid (FA) where the last four elution steps included a 5 min incubation at 95°C prior to centrifugation. Eluted peptides were StageTip purified and analyzed by mass-spectrometry.

The LC-MS analysis was conducted using an EASY-nLC 1200 UHPLC system (Thermo Fisher) connected to an Orbitrap Lumos mass spectrometer (Thermo Fisher). The nLC system utilized a single analytical column set up, employing PicoFrit Emitters (New Objectives, 75 μm inner diameter) that were packed in-house with Reprosil-Pure-AQ C18 material (Dr. Maisch, 1.9-μm particle size, 19-21 cm column length) and was operated with the flow rate 200 nl/min. All samples dissolved in 0.1% formic acid were injected onto the column and eluted in a gradient from 2 to 25% acetonitrile for 95 min and from 25% to 80% acetonitrile for 10 min, followed by isocratic elution at 80% acetonitrile for 15 min (total elution time 120 min). The nanoSpray ion source was operated at 2.1 kV spray voltage and 300°C heated capillary temperature. A precursor MS1 scan (m/z 350– 1,700) of intact peptides was performed in the Orbitrap at a resolution of 120,000. This was followed by high-energy collision dissociation (HCD)-MS2 with the collision energy for HCD scans was set to 37% +/- 5% at a resolution of 60,000 for the fifteen most abundant multiply charged precursors in the MS1 spectrum. A minimum MS1 signal threshold of 50,000 was set to trigger data-dependent fragmentation. A dynamic exclusion window of 30 seconds was used to avoid repeated analysis of the same species. The MS/MS spectra were processed against the non-redundant human proteome using the SEQUEST-HT search engine in Proteome Discoverer 2.2. The search parameters were defined as follows: semi enzymatic tryptic cleavage with up to 2 missed cleavages, variable methionine oxidation, N-Term Biotin, lysine Biotin, N-Term TMT6plex and lysine TMT6plex. Fixed modification was set for cysteine carbamidomethylation. The initial search was performed on a 10ppm precursor mass tolerance and 0.02 Da fragment ion tolerance with a false discovery rate (FDR) of 1%.

### Production and purification of soluble LRP1 ectodomain

HEK-sLRP1 and HEK-sLPR1^ΔT11^ were cultured in suspension in Freestyle F17 Expression Medium (Gibco™, Thermo Fisher) supplemented with 1% FBS, 0.1% Kolliphor P188 (Sigma) and 2 mM GlutaMax at 37°C and 5% CO_2_ under constant agitation (180 rpm). Clarified culture supernatant was depleted of free biotin using BioLock (IBA LifeSciences 2-0205-050) and made 25 mM Tris pH 8, 150 mM NaCl, 2 mM CaCl_2_, 1 mM DTT (Buffer A), and loaded on a Strep-Tactin®XT 4Flow® (IBA Lifesciences 2-5010-010) affinity column equilibrated in Buffer A. The column was washed with 25 mM MES pH 5.5, 150 mM NaCl, 1 mM DTT followed by Buffer A, and eluted with 50 mM Biotin in Buffer A. The LRP1 ectodomain was further purified by size-exclusion chromatography on a Superose 6 Increase 10/300 GL column (Cytiva) equilibrated with 25 mm MES pH 5.5, 150 mm NaCl, 4 mm CaCl_2_, 1 mM DTT. LRP1 containing fractions were pooled and passed over a Zeba Spin column equilibrated in 25 mM HEPES pH 7.5, 150 mM NaCl, 4 mM CaCl_2_+, 1 mM TCEP.

### Released O-Glycan analysis

Purified sLRP1, sLPR1^ΔT11^ and a similarly produced sMUC1 control sample were dissolved in 50 mM Tris HCl, 100 mM NaCl and 1× cOmplete™ protease inhibitor (EDTA-free; Roche) with a protein concentration of 0.4 µg/µL. From each sample, 25 µL was blotted on a PVDF membrane in filter plate format and N-glycans were removed using PNGase F as described previously^39,75^. O-Glycans were released using a non-reductive beta-elimination approach using 20% 1,8-diazabicyclo(5.4.0)undec-7-ene and 20% hydroxylamine for 1 h at 37°C, and subsequently labeled with 2-aminobenzamide (2-AB), HILIC and PGC SPE purified and resuspended in 20 µL water for LC-MS analysis^39^. Two microliter per sample (10% of total) was injected per analysis. The glycans were separated by nano-flow liquid chromatography (nanoLC) using a single analytical column setup packed with Reprosil-Pure-AQ C18 phase (Dr. Maisch, 1.9 μm particle size, 19–21 cm column length) in an EASY-nLC 1200 UHPLC (Thermo Fisher) using a PicoFrit Emitter (New Objectives, 75 μm inner diameter). The emitter was interfaced to an Orbitrap Fusion Lumos MS (Thermo Fisher) via a nanoSpray Flex ion source. An 1 h method was used with a gradient from 3% to 32% of solvent B in 35 min, from 32% to 100% B in the next 10 min and 100% B for the last 15 min at 200 nL/min (solvent A: 0.1% formic acid in water; solvent B: 0.1% formic acid in 80% ACN). A precursor MS scan (*m/z* 200-1700, positive polarity) was acquired in the Orbitrap at a nominal resolution of 120,000, followed by Orbitrap higher-energy C-trap dissociation (HCD)-MS/MS at a nominal resolution of 50,000 of the 10 most abundant precursors in the MS spectrum (charge states 1 to 4). A minimum MS signal threshold of 30,000 was used to trigger data-dependent fragmentation events. HCD was performed with an energy of 27% ± 5%, applying a 20 s dynamic exclusion window. MS1 feature detection in the raw files was performed using Thermo Proteome Discoverer 2.2.0.388 (Thermo Fisher). The identified LC-MS features (defined by *m/z*, retention time (RT) and charge) were filtered based on RT (> 10 min, < 35 min) and charge (≥ 1, ≤ 3). The [M+H] values of the resulting features were imported into GlycoWorkbench 2.1 (build 146) and matched to glycan compositions with 0 to 8 hexoses, 0 to 8 *N*-acetyl hexosamines, 0 to 3 fucoses, 0 to 4 *N*-acetyl neuraminic acids and a 2-AB label. An additional matching was performed to glycan compositions with 0 to 6 hexoses, 0 to 6 *N*-acetyl hexosamines, 0 to 2 fucoses, 0 to 2 *N*-acetyl neuraminic acids, 0 to 3 pentoses and a 2-AB label. The complete list of identified compositions was imported into Skyline 21.1.0.146 (ProteoWizard), using the Molecule Interface. Extracted ion chromatograms were generated for the first three isotopologues of each glycan. Chromatographic peaks were manually selected based on accurate mass (> -1 ppm, < 1 ppm), isotopic dot product (idotp; > 0.85) and a minimal signal intensity of 1×10^6^ in at least one of the samples and integrated for all samples. MS/MS spectra were manually assigned for each MS1 feature in at least one sample. All MS1 signals were integrated and normalized to the total sum of glycans per sample, or alternatively against the total intensity sum of all *O*-GalNAc glycans per sample. Glycans were grouped on structural features and relative abundances of core1, core2 and sialylation were calculated.

### Surface plasmon resonance spectroscopy

Surface plasmon resonance (SPR) spectroscopy was performed on a Biacore 3000 instrument (Cytiva), running at 25°C and a data collection rate of 1 Hz. The running buffer was 10 mm HEPES, pH 7.5, 150 mM NaCl, 2 mM CaCl_2_ and 0.05% Tween 20. Ligands were immobilized on CM5 chips according to the instructions of the manufacturer (Cytiva). In short, surfaces were activated by a 7 min injection of a 1:1 mixture of 0.1 M NHS (N-hydroxysuccinimide) and 0.4 M EDC (3-(N,N-dimethylamino) propyl-N-ethylcarbodiimide). sLRP1 and sLPR1^ΔT11^ diluted to 10 µg/ml in 10 mM sodium acetate pH 4.5, were then injected over the activated surfaces until a density of 5,000 RU had been reached. Residual reactive groups were blocked by a 7 min injection of 1 M ethanolamine pH 8.5. For determination of steady state kinetics, a two-fold titration series of RAP ranging from 1-256 nM was injected over the surfaces for 300 s followed by a 120 s dissociation phase. At the end of each cycle, surfaces were regenerated by a 60 s injection of 1 M ethanolamine pH 8.5. Using the BIAevaluation 4.1.1 software (Cytiva), recorded signals from the active flow cells were double referenced, i.e. the signal from the in-line reference flow cell was subtracted as was the signal from a blank run (0 nM analyte). Steady state K_D_ values were determined using GraphPad Prism version 10.1 and the built-in “Two sites – specific binding” equation.

### Mass photometry

Microscope glass slides (High precision 24 x 50 mm No.1.5 coverslips, Thorlabs cat. CG15KH) were cleaned by sequential rinses of isopropanol (HPLC grade) and Milli-Q H_2_O for 5 cleaning cycles, followed by drying with a clean nitrogen stream. Four gaskets (Reusable culturewell™ gaskets 3 mm diam. × 1 mm depth, cat. GBL103250-10 EA, Sigma-Aldrich) were cut to 2 × 2 array, cleaned similarly to the glass slides. A drop of immersion oil (Thorlabs MOIL-30) was placed on the mass photometry (OneMP, Refeyn, Ltd, Oxford, UK) objective and the glass slide and gasket adhered to the glass slide was placed on top of the immersion oil and positioned on the instruments mobile stage. To focus the objective, 15 µl of fresh buffer was added to the well, and the focal position was identified and secured in place with the autofocus system based on total internal reflection for the entire measurement. To calibrate the mass photometry instrument following autofocus stabilization, 5 μl of proteins standards of molecular weights 66 kDa, 146 kDa, 242 kDa and 480 kDa from native mark unstained protein standard (Thermo Fisher LC0725) was added into the well, mixed by pipetting up and down three times and movies of 120 s duration were recorded. Immediately prior to mass photometry measurements, protein stocks were diluted in HEPES pH 7.5, 150 mM NaCl, 4 mm CaCl2, 1 mM TCEP. The same procedure was followed for each sample, where 15 µl of buffer was added to the well, autofocused, and subsequently 5 µl of diluted protein (nanomolar concentrations) was added to individual wells. Each sample was measured at least three times independently (n ≥ 3). Data was acquired using an OneMP mass photometer (Refeyn Ltd, Oxford, UK). Data acquisition was performed using AcquireMP (Refeyn Ltd, v2.2) and data analysis and graphing was performed in R.

### Computational Methods

The 3D structure of the non-glycosylated CR5-CR6 in complex with RAP domain1 (RAPd1) obtained from NMR-guided molecular docking (PDB 2fyl) was chosen as starting structure for all MD simulations. Core 1 and sialylated core 1 O-glycans were added at Thr44 to this structure based on a scheme now automated in GlycoShape^76^, where the glycan 3D structures from the GlycoShape database are linked to the protein with the tool Re-Glyco. We ran to completion three independent MD simulations for the complex with a core1-glycosylated linker and two simulations for the complex sialyl-core1 glycosylated linker. We collected 1 µs of data from each simulation, for a total of 5 µs of cumulative sampling. In all cases, we used the same MD setup protocol, which started with a geometry optimization of 500k steps of steepest descent energy minimization. The system was then heated in two steps in the NVT ensemble, from 0 K to 100 K over 500 ps, and then from 100 K to 300 K over another 500 ps interval, using Langevin dynamics with a friction coefficient of 1.0 ps⁻¹. A 500 ps equilibration phase in the NPT ensemble was then ran to bring the system up to 1 atm of pressure using a Berendsen barostat. All MD simulations were run using version 2018 of the AMBER package on GPUs^77^. All bonds to hydrogen atoms were restrained using the SHAKE algorithm to allow for an integration time step of 2 fs. Long-range electrostatic interactions were treated using the Particle Mesh Ewald (PME) and an 11 Å cutoff was used for both electrostatic and van der Waals interactions. The GLYCAM_06j-1 force field^78^ was used to represent carbohydrate atoms, the ff14SB parameters^79^ for the protein and counterions, and the TIP3P model^80^ was used to represent water molecules.

### Statistical analysis

Differences in ligand uptake as measured by flow cytometry were assessed using the ratio paired *t*-test in GraphPad Prism version 10.2.3. The significant accumulation (*p* < 0.05) of biotinylated CSF proteins were determined using a two-tailed unpaired *t*-test and computed in R.

## Data availability

The data supporting the findings of this study are available within the paper and its Supplementary Information files, and available from the corresponding author upon request. The mass spectrometry proteomics data have been deposited to the ProteomeXchange Consortium via the PRIDE^81^ partner repository with the dataset identifier PXD058370.

## Acknowledgments

This work was primarily supported by a Novo Nordisk Foundation Hallas Møller Ascending Investigator grant no NNF0073793 and a Sapere Aude Research Leader grant from the Independent Research Fund Denmark grant no. 2066-00043B (K.T.S.). We acknowledge the National Institute of Health, NIH grant no R01 AG073236 (B.H. and D.K.S.), the JPB and Rainwater Foundation (B.H.), Cure Alzheimer Fund (B.H.), the Independent Research Fund Denmark grant no. 1056-00022B (J.H), Novo Nordisk Foundation grant no NNF0067602 (T.D.M.), Grupos Consolidados Gobierno Vasco 2021 grant no. 449IT1720-22 (C.M.), Proyectos de Generación de Conocimiento from the Ministerio de Ciencia, Innovación y Universidades grant no. PID2022-136788OB-I00 (C.M.). The Science Foundation of Ireland (SFI) Frontiers for the Future Programme is gratefully acknowledged for financial support of Silvia D’Andrea’s postgraduate training (20/FFP-P/8809). The Irish Centre for High-End Computing (ICHEC) is gratefully acknowledged for generous allocation of computational resources. We thank Jakob Hauge Mikkelsen (Biacore – Molecular Interactions Core Facility, Aarhus University) for SPR analysis, Christina Christoffersen (University of Copenhagen, Department of Biomedical Sciences) for providing CSF samples, and Claire B. Holden for excellent technical assistance.

## Author Contributions

This research was conceptualized by K.T.S and J.H., executed by J.H., A.B.H., T.M.D., S.J., S.A., N.H., L.H.H., R.B., N.Q., Z.Y., S.Y.V., R.L.M., supervised by C.M., D.K.S., B.H, E.F. and K.T.S, and written by J.H. and K.T.S.

## Competing interests

The authors report no conflicts of interest.

**Correspondence** and requests for materials should be addressed to Katrine T. Schjoldager.

## Notes

### Competing Interest Statement

The authors have declared no competing interest.

