## Supplementary Information for "The Ligand Preference of LRP1 is Regulated by O-glycans"

**This PDF file includes:**

- Supplementary Figures 1-8
- Supplementary Tables 1-4

**a**

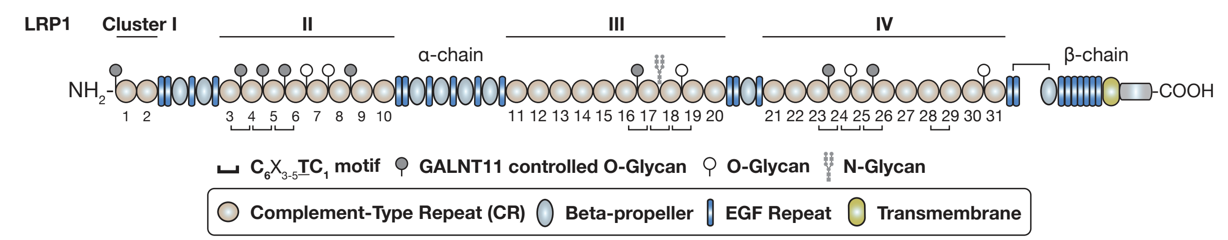


**b**

**Cluster I**

| | CR1–CR2 | CR2– |
|---|---|---|
| Human | DAPKTC | CPQSKAQRC |
| Mouse | DAPKTC | CPQSKAQRC |
| Chicken | EAPKTC | CPQSKVSRC |
| Zebrafish | EAPKTC | CHHTQVTQC |
| Lamprey | -APKTC | CQKPNEMKC |
| Conservation | :**** | : : : |

**Cluster II**

| | CR3–CR4 | CR4–CR5 | CR5–CR6 | CR6–CR7 | CR7–CR8 | CR8–CR9 | CR9–CR10 |
|---|---|---|---|---|---|---|---|
| Human | CHQHTC | CSARTC | CAYPTC | CSHSC | CTNQATRPPGGC | CEGVTHVC | CESLAC |
| Mouse | CHQHTC | CSARTC | CAYPTC | CSHSC | CTNQATRPPGGC | CEGVTHVC | CEALAC |
| Chicken | CHQHTC | CSARTC | CAYPTC | CSHSC | CTNQATRPPGGC | CEGVTHVC | CESLVC |
| Zebrafish | CNQHTC | CLARTC | CAYPTC | CSHSC | CTNQATRPPGGC | CEGVTHMC | CEALEC |
| Lamprey | CHQHTC | CAARTC | CAYPTC | CNHSC | CTKLATRPPGGC | CDGVVRTC | CETMLC |
| Conservation | :*** | *** | **** | .** | *: ******* | ****** | *:: |

**Cluster III**

| | CR11–CR12 | CR12–CR13 | CR13–CR14 | CR14–CR15 | CR15–CR16 | CR16–CR17 | CR17–CR18 | CR18–CR19 | CR19–CR20 |
|---|---|---|---|---|---|---|---|---|---|
| Human | CNSRRC | CNKTAC | CSATDC | CPGV-KRPRC | C----NKFC | CEGKTC | CLYNSTC | CEYPTC | CTSQEHKC |
| Mouse | CNSRRC | CNKTAC | CSATDC | CPGV-KRPRC | C----NKFC | CEGKTC | CLYNSTC | CEYPTC | CTSPEHKC |
| Chicken | CSSRKC | CNKTSC | CTATDC | CPGG-RKPKC | CSERQDKFC | CRLTTC | CLYNNTC | CEYPTC | CSSPESKC |
| Zebrafish | CANRMC | CNMTLC | CSPTDC | CPEK-RQQKC | CEKF-FQFC | CKAKTC | CVFNKTC | CEYPTC | CTDTESRC |
| Lamprey | CTSRQC | CNNSTC | CGPMNC | CKKKSGSPGC | C----DMLC | CTFRAC |  |  | CSANNSKC |
| Conservation | .* | * : | : | | . | : | ***** | **** | : : : |

**Cluster IV**

| | CR21–CR22 | CR22–CR23 | CR23–CR24 | CR24–CR25 | CR25–CR26 | CR26–CR27 | CR27–CR28 | CR28–CR29 | CR29–CR30 | CR30–CR31 |
|---|---|---|---|---|---|---|---|---|---|---|
| Human | CPEFKC | CDIHVC | CPEVTC | CTQMTC | CDERTC | CTPRPC | CTPRC | CG-TGVRTC | CARFVC | CEPPTAHTTHC |
| Mouse | CPEFKC | CDIHVC | CPEVTC | CTQMTC | CDERTC | CTPRPC | CTPRC | CG-TGVRTC | CARFIC | CEPPTAQNPHC |
| Chicken | CPEFKC | CDIHVC | CPEVTC | CTQMTC | CDERTC | CTPRPC | CIPRC | CG-TGVRTC | CLKFQC | CESPTAKPKSC |
| Zebrafish | CPEFKC | CDIHVC | CPEVTC | CTQMKC | CDERTC | CVPRQC | CDLKC | CEITEVRHC | CRHFQC | CDYES----TC |
| Lamprey | CPDFKC | CDVHVC | CPEVTC | CTQMTC | CEDRTC | CTPRPC | CSPRC | CA-AIVRTC | CLKFQC | CEKPSPKPRIC |
| Conservation | *:** | *:** | **** | ***. | ::** | .** | : | : ** | :* | : : |

**Supplementary Fig. 1 | O-glycosylation of LRP1 CR linkers. a** Same as in Fig. 1a. Schematic representation of LRP1 displaying the various domains. Indicated are sites of Complement-Type repeat (CR) linker O-glycosylation identified by O-glycoproteomic analyses of various mammalian tissues and cell lines. These analyses include samples with disrupted GALNT11 gene. **b** Alignment of LRP1 linker sequences across different animal species. The complete LRP1 sequences were aligned with the T-Coffee multiple sequence alignment tool using default settings. LRP1 Uniprot IDs: Human Q07954, Mouse Q91ZX7, Chicken P98157, Zebrafish F1QY34, Lamprey S4RNX1.

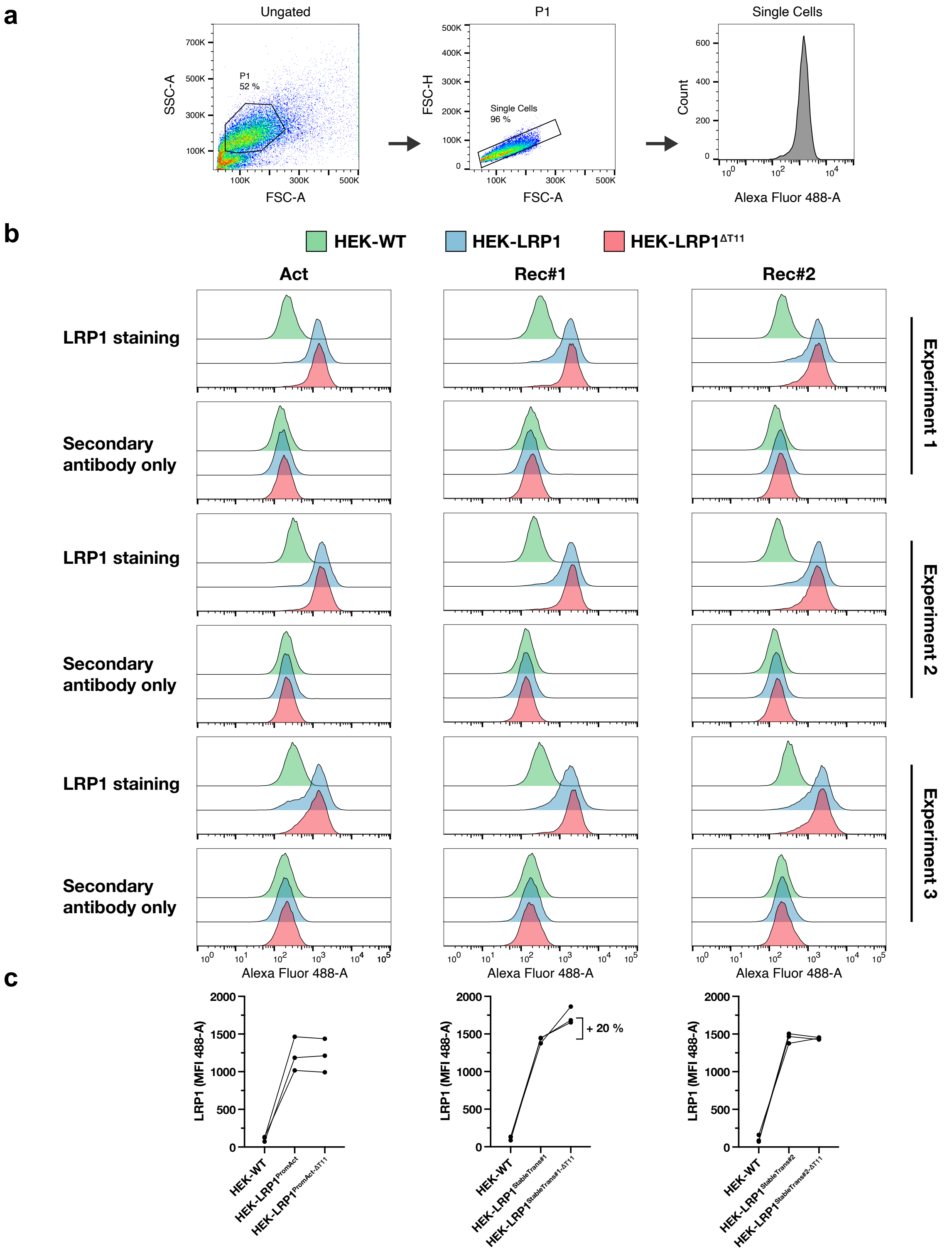


**Supplementary Fig. 2 | Flow cytometric measurement of LRP1 surface expression in HEK-LRP1 cells.** Live cells were stained with anti-LRP1 mAb 8G1 followed by Alexa Fluor 488 conjugated secondary antibody. **a** The gating strategy used to measure Alexa Fluor 488-A signal from singlet cells. **b** Histogram overlays of LRP1 cell surface expression from three independent experiments for each HEK-LRP1 cell line. **c** Background subtracted LRP1 median fluorescence intensities (MFIs) for the experiments shown in b.

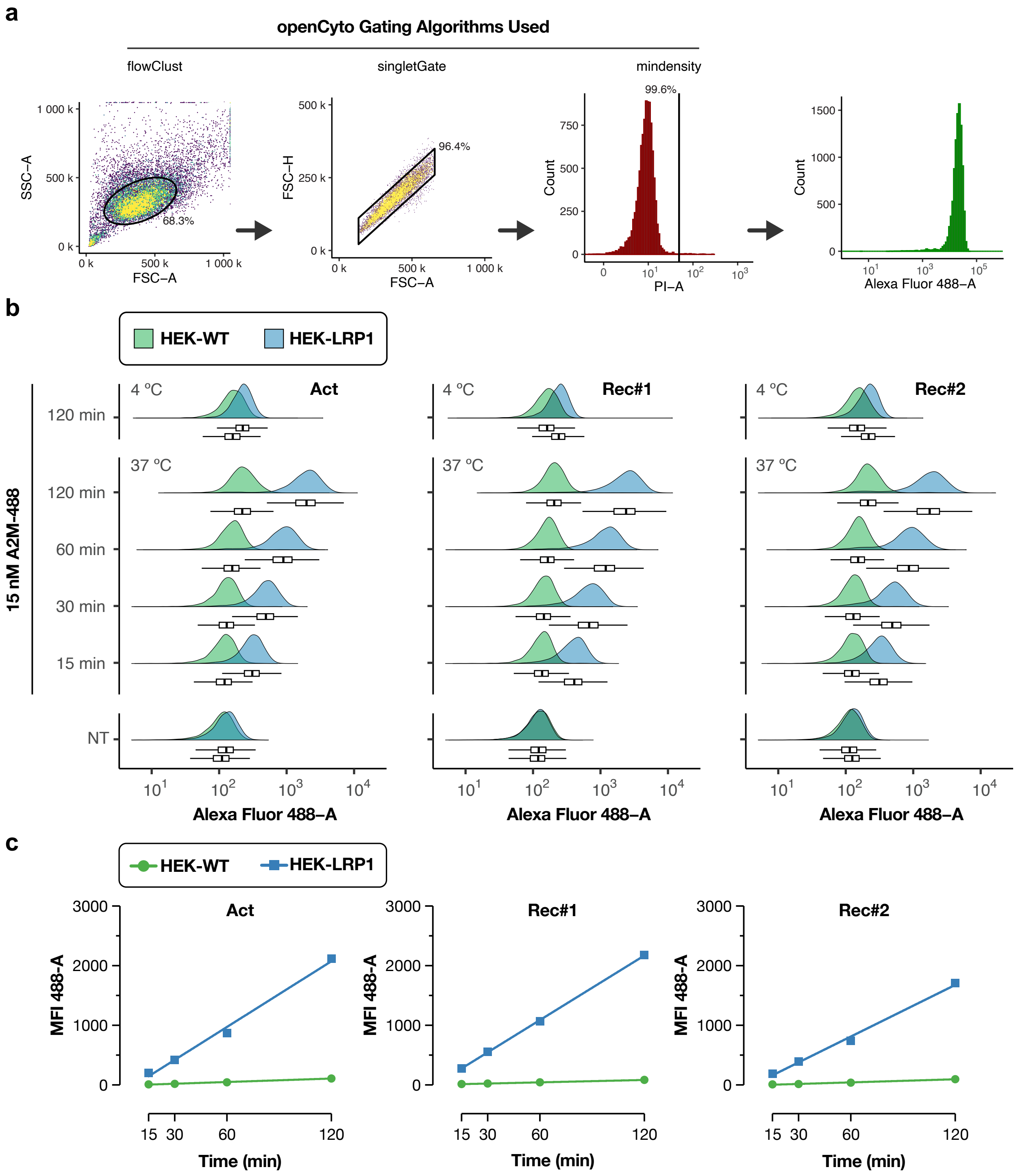


**Supplementary Fig. 3 | Time-dependent A2M uptake in HEK-LRP1 cells.** Cells cultured in 96-well plates were incubated at 37°C with 15 nM of Alexa Fluor 488-labeled Alpha-2-Macroglobulin (A2M) diluted in complete growth medium for indicated times before being released, stained with propidium iodide (PI) and analyzed by flow cytometry. **a** Gating strategy to record 488 fluorescence from single-cell and PI-negative gated cells. This gating strategy applies to Fig. 2, Fig. 3d, and Supplementary Fig. 3-4**. b** Histogram overlays with corresponding Box-and-Whisker plot in the style of Tukey of recorded 488-A signal. **c** Incubation time at 37°C plotted against background subtracted 488-A MFI values.

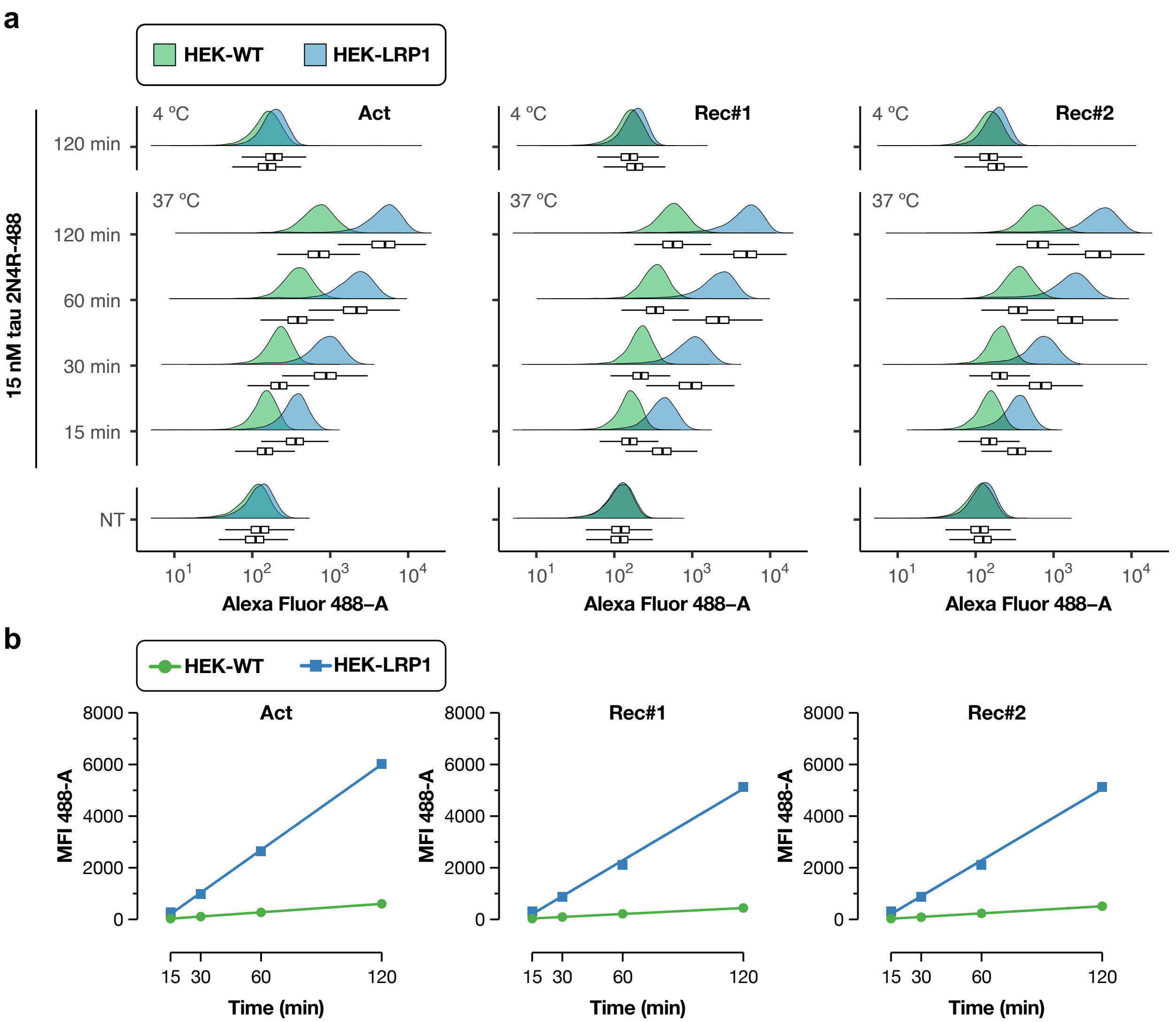


**Supplementary Fig. 4 | Time-dependent tau uptake in HEK-LRP1 cells.** Cells cultured in 96-well plates were incubated at 37°C with 15 nM of Alexa Fluor 488-labeled tau 2N4R diluted in complete growth medium for indicated times before being released, stained with propidium iodide (PI) and analyzed by flow cytometry. **a** Histogram overlays with corresponding Box-and-Whisker plot in the style of Tukey of recorded 488-A signal. **b** Incubation time at 37°C plotted against background subtracted 488-A MFI values.

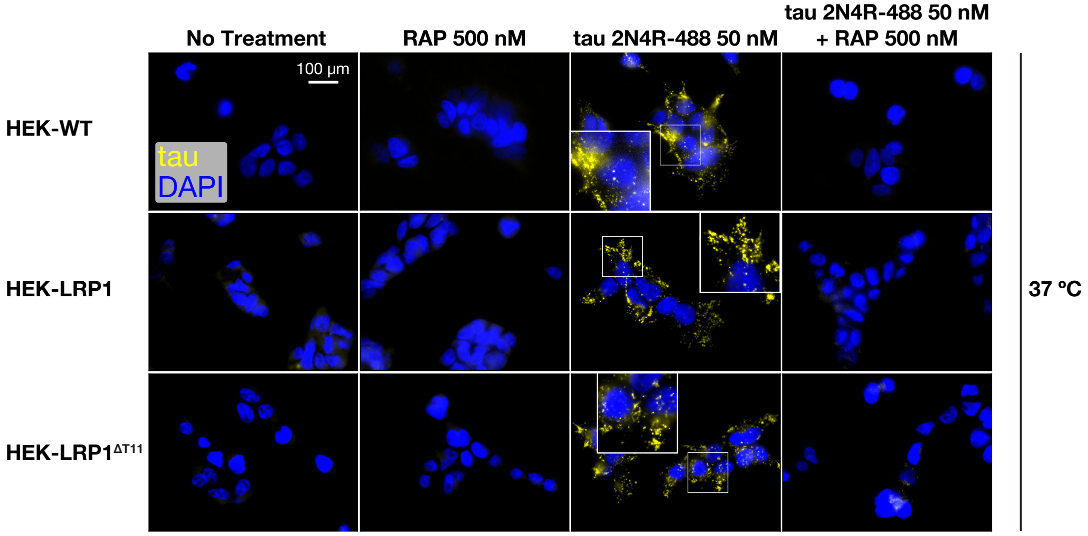


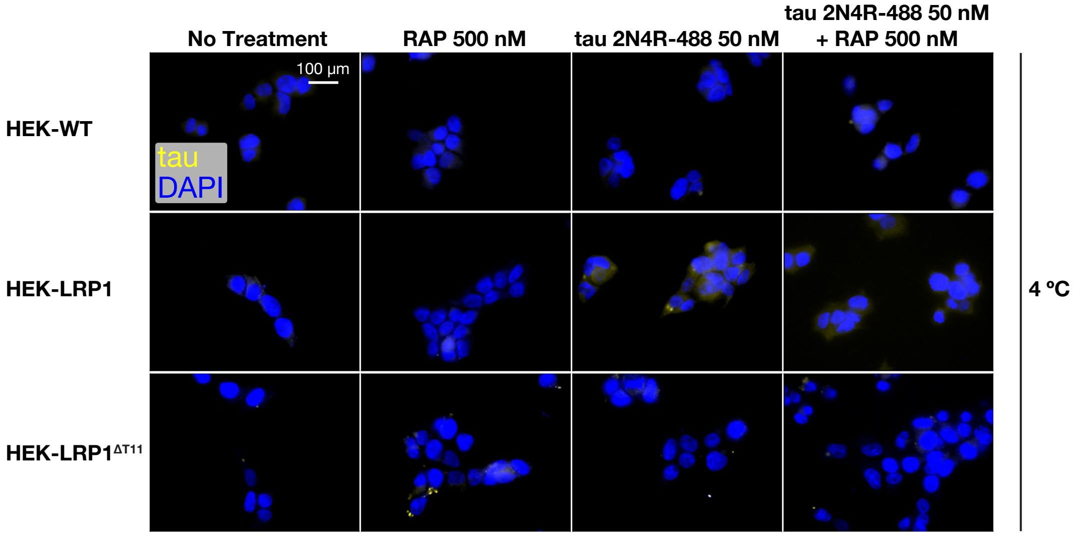


**Supplementary Fig. 5 | Visualization of tau uptake in HEK-LRP1 cells.** Cells cultured on coverslips were incubated for 2 hours at 37°C or 4°C with 50 nM tau 2N4R-488 (+/- 500 nM RAP) diluted in complete growth medium before being washed and fixed. The coverslips were mounted with ProLong™ Gold Antifade supplemented with DAPI (Thermo Fisher P36931).

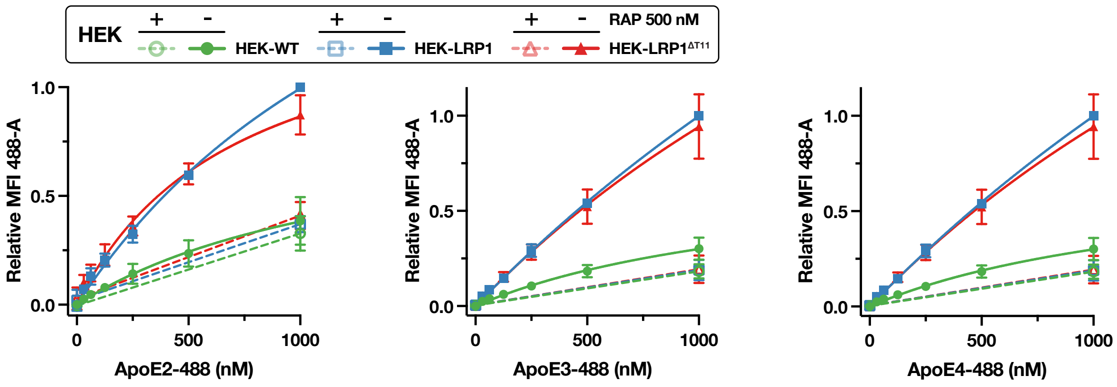


**Supplementary Fig. 6 | Uptake of ApoE in HEK-LRP1 cells.** Cells were incubated at 37°C for 2 hours with fluorescently labeled ApoE proteins diluted in Opti-MEM medium before being released, stained with propidium iodide (PI) and analyzed by flow cytometry. Each graph represents the aggregate of three experiments, one for each of the HEK-LRP1$^{Act}$, HEK-LRP1$^{Rec\#1}$ and HEK-LRP1$^{Rec\#2}$ lines. Data are presented as mean ± SD.

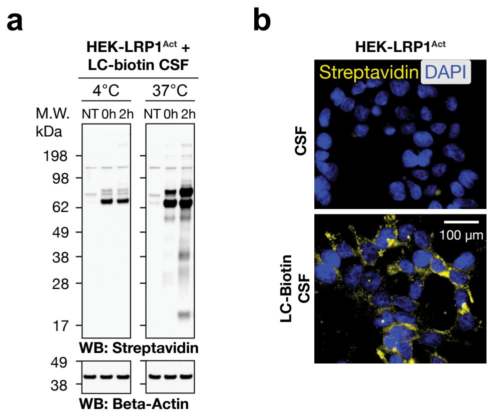


**Supplementary Fig. 7 | Uptake of biotinylated CSF proteins. a** Cells were cultured in the presence of CSF proteins labeled with LC-Biotin for up to 2 hours at indicated temperatures before being washed and harvested. Total cell lysates were analyzed by streptavidin western blot. **b** Cells were cultured on coverslips for 2 hours in CSF or biotinylated CSF before being washed, fixed with paraformaldehyde and stained with streptavidin Alexa Fluor 488 conjugate. CSF, cerebrospinal fluid; NT, no treatment.

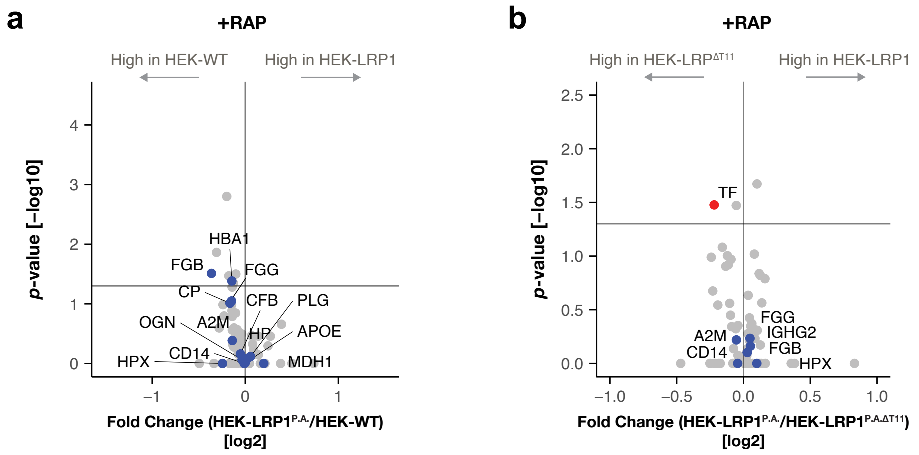


**Supplementary Fig. 8 | RAP inhibition in mass-spectrometric quantification of endocytosed LC-Biotin tagged CSF proteins.** Related to Fig. 3. Volcano plots of those LC-biotin-labeled proteins identified when cells were co-incubated with LC-biotin labeled CSF proteins and 500 nM RAP with comparison of (**a**) HEK-LRP1$^{Act}$ to HEK-WT and (**b**) HEK-LRP1$^{Act}$ to HEK-LRP1$^{Act\text{-}\Delta T11}$. Those LC-biotin-labeled proteins found significantly endocytosed ($p$ < 0.05) in the absence of RAP (Fig. 3b, c) are colored blue and annotated.

**Supplementary Table 1 |** Biotinylated CSF proteins accumulating in HEK-LRP1$^{Act}$ over HEK-WT cells (*p* < 0.05). Related to Fig. 3b.

| Uniprot ID | Protein Name | Gene Name | log2 fold change (HEK-LRP1$^{Act}$ /HEK-WT) | Linear fold change (HEK-LRP1$^{Act}$ /HEK-WT) |
|---|---|---|---|---|
| P01023 | Alpha-2-macroglobulin | A2M | 1.29 | 2.45 |
| P02790 | Hemopexin | HPX | 1.15 | 2.22 |
| P20774 | Mimecan | OGN | 1.01 | 2.01 |
| P69905 | Hemoglobin subunit alpha | HBA1 | 0.76 | 1.69 |
| P02675 | Fibrinogen beta chain | FGB | 0.72 | 1.65 |
| O14498 | Immunoglobulin superfamily containing leucine-rich repeat protein | ISLR | 0.71 | 1.63 |
| P02679 | Fibrinogen gamma chain | FGG | 0.56 | 1.47 |
| P07225 | Vitamin K-dependent protein S | PROS1 | 0.50 | 1.42 |
| P40925 | Malate dehydrogenase, cytoplasmic | MDH1 | 0.36 | 1.28 |
| P00450 | Ceruloplasmin | CP | 0.29 | 1.22 |
| P00751 | Complement factor B | CFB | 0.29 | 1.22 |
| P00738 | Haptoglobin | HP | 0.29 | 1.22 |
| P00747 | Plasminogen | PLG | 0.23 | 1.17 |
| P08571 | Monocyte differentiation antigen CD14 | CD14 | 0.17 | 1.12 |
| P02649 | Apolipoprotein E | APOE | 0.12 | 1.09 |

**Supplementary Table 2 |** Biotinylated CSF proteins accumulating in HEK-LRP1$^{Act}$ over HEK-LRP1$^{Act\text{-}\Delta T11}$ cells ($p$ < 0.05). Related to Fig. 3c.

| Uniprot ID | Protein Name | Gene Name | log2 fold change (HEK-LRP1$^{Act}$ /HEK-LRP1$^{Act\text{-}\Delta T11}$) | Linear fold change (HEK-LRP1$^{Act}$ /HEK-LRP1$^{Act\text{-}\Delta T11}$) |
|---|---|---|---|---|
| P02790 | Hemopexin | HPX | 0.73 | 1.66 |
| P02675 | Fibrinogen beta chain | FGB | 0.28 | 1.21 |
| P01023 | Alpha-2-macroglobulin | A2M | 0.24 | 1.18 |
| O94769 | Extracellular matrix protein 2 | ECM2 | 0.20 | 1.15 |
| P02679 | Fibrinogen gamma chain | FGG | 0.18 | 1.13 |
| P01859 | Immunoglobulin heavy constant gamma 2 | IGHG2 | 0.13 | 1.09 |
| P08571 | Monocyte differentiation antigen CD14 | CD14 | 0.06 | 1.04 |

**Supplementary Table 3 |** Antibodies, lectins and streptavidin conjugates used in this study.

| Reagent | Species; Clonality | Source; Catalog No | Application | Dilution; Working Concentration (ug/mL) |
|---|---|---|---|---|
| anti-LRP1 α-chain 8G1 | Mouse; Monoclonal | Abcam; ab20384 | WB/Flow/IF | 1:400; 5 |
| anti-Beta Actin | Mouse; Monoclonal | Proteintech; 66009-1-Ig | WB | 1:20,000; 0.05 |
| anti-Mouse HRP | Rabbit; Polyclonal | Agilent; P0260 | WB | 1:4,000 |
| anti-Mouse IgG Alexa Fluor 647 | Goat; Polyclonal | Thermo; A-21236 | Flow/IF | 1:1,000; 2 |
| anti-Mouse IgG Alexa Fluor 488 | Goat; Polyclonal | Thermo; A-11001 | Flow/IF | 1:1,000; 2 |
| Jacalin Lectin Biotinylated | NA | Vector labs; B-1155-5 | LB | 1:10,000; 0.5 |
| Streptavidin HRP | NA | Agilent; P039701-2 | WB | 1:4,000 |
| Streptavidin Alexa Fluor 488 | NA | Thermo; S11223 | IF | 1:2,000: 1 |

**Supplementary Table 4 |** Sequencing and insertion/deletion analysis of *GALNT11* KO clones.

| **Cell line** | ***GALNT11* exon 2 Insertions and Deletions** |
|---|---|
| HEK-LRP1$^{Act\text{-}\Delta T11}$ | WT :CAGTGACC--GCTtggGCTACCACAG<br>+1 :CAGTGACCC-GCTtggGCTACCACAG<br>+2 :CAGTGACCCCGCTtggGCTACCACAG |
| HEK-LRP1$^{Rec\#1\text{-}\Delta T11}$ | WT :CAGTGACC-GCTtggGCTACCACAG<br>+1 :CAGTGACCCGCTtggGCTACCACAG |
| HEK-LRP1$^{Rec\#2\text{-}\Delta T11}$ | WT :CAGTGACC-GCTtggGCTACCACAG<br>+1 :CAGTGACCCGCTtggGCTACCACAG |
| SH-SY5Y$^{\Delta T11,\ 2D5}$ | WT :CAGTGACC-GCTtggGCTACCACAG<br>+1 :CAGTGACCCGCTtggGCTACCACAG<br>-2 :CAGTGACC---TtggGCTACCACAG |
| SH-SY5Y$^{\Delta T11,\ 2C7}$ | WT :CAGTGACC-GCTtggGCTACCACAG<br>+1 :CAGTGACCCGCTtggGCTACCACAG |
| SH-SY5Y$^{\Delta T11,\ 2C8}$ | WT :CAGTGACC-GCTtggGCTACCACAG<br>+1 :CAGTGACCCGCTtggGCTACCACAG<br>-1 :CAGTGAC--GCTtggGCTACCACAG |
| HEK$^{KI\ RAP}$-sLRP1$^{\Delta T11}$ | WT :CAGTGACC----GCTtggGCTACCACAG<br>+1 :CAGTGACCC---GCTtggGCTACCACAG<br>+4 :CAGTGACCTCACGCTtggGCTACCACAG |
